# Multi-omic profiling of pathogen-stimulated primary immune cells

**DOI:** 10.1101/2023.09.13.557514

**Authors:** Renee Salz, Emil E. Vorsteveld, Caspar I. van der Made, Simone Kersten, Merel Stemerdink, Tabea V. Riepe, Tsung-han Hsieh, Musa Mhlanga, Mihai G. Netea, Pieter-Jan Volders, Alexander Hoischen, Peter A.C. ’t Hoen

## Abstract

We performed long-read transcriptome and proteome profiling of pathogen-stimulated peripheral blood mononuclear cells (PBMCs) from healthy donors to discover new transcript and protein isoforms expressed during immune responses to diverse pathogens. Long-read transcriptome profiling reveals novel sequences and isoform switching induced upon pathogen stimulation, including transcripts that are difficult to detect using traditional short-read sequencing. Widespread loss of intron retention occurs as a common result of all pathogen stimulations. We highlight novel transcripts of *NFKB1 and CASP1* that may indicate novel immunological mechanisms. RNA expression differences did not result in differences in the amounts of secreted proteins. Clustering analysis of secreted proteins revealed a correlation between chemokine (receptor) expression on the RNA and protein levels in *C. albicans-* and Poly(I:C)-stimulated PBMCs. Isoform aware long-read sequencing of pathogen-stimulated immune cells highlights the potential of these methods to identify novel transcripts, revealing a more complex transcriptome landscape than previously appreciated.

## Introduction

Immune system responses within the context of an infection are shaped by the nature of the infection and by inter-individual variability, contributing to differential susceptibility to infections and to various diseases with an inflammatory component. Dynamic expression of transcripts and proteins in a range of cells responsible for the innate immune response is important to shape the first line of defense against a wide variety of pathogens^1^. Pattern recognition receptors (PRR) initiate acute inflammatory responses, activating signaling cascades that converge on various transcription factors. Multiple levels of regulation orchestrate the dynamic expression of transcripts and proteins, including transcriptional and post-transcriptional checkpoints such as mRNA splicing and protein translation. Examples of this include the regulatory role of alternative splicing of Toll-like receptors (TLR) and their downstream signaling factors^2, 3^.

Methods for investigating innate immune responses include *in vitro* stimulation of primary immune cells with pathogens or microbial components. These methods allow for specific investigation of host-pathogen interactions that shape the immune response elicited by specific cell types and have been under extensive investigation in research on the innate immune system^4, 5^. Stimuli that are commonly used include molecules that stimulate a specific TLR, such as *E. coli* lipopolysaccharide (LPS) for TLR4^6^, dsRNA mimicking Poly(I:C) for TLR3^7^ and imidazoquinolines for TLR7/8^8^. Stimulation is also often elicited by live or heat-killed pathogens^9, 10^, which stimulate at the same time a broader range of PRRs^11, 12^.

Transcriptome characterization is traditionally performed using short read RNA sequencing. These sequencing approaches are limited by their short-read length (approximately 150-300 bp), necessitating the computational reconstruction of whole transcripts and making detection of different transcripts of the same gene inaccurate. This limitation is especially pronounced in immune biology, where tight regulation of isoform expression has previously been described to play a major role in processes, such as the expression of multiple IL-32 transcripts with different inflammatory potency^13^ and alternative splicing of *CD45* in T cell activation^14^. Recent long-read sequencing approaches provide a more complete and accurate reflection of the transcriptome. Sequencing technologies provided by PacBio and Oxford Nanopore allow for the sequencing of mRNA (or cDNA) molecules from the ultimate 3’-end to the ultimate 5’-end, which have given a more comprehensive view into the complexity of the transcriptome. A number of studies have indicated that the isoform landscape is much more complex than previously appreciated^15–17^. Long-read mRNA sequencing has provided insight into regulatory mechanisms of immune responses, for instance in alternative splicing in macrophages^18^, and allows for accurate sequencing of complex transcripts in immune cells^19^. Further work on cell-type specific long-read transcriptomes have shown preferential expression of transcripts in specific cell types^20^.

The impact of the newly discovered transcripts as well as post-transcriptional processes can only be fully understood by observing the proteome. Many studies have characterized the transcriptomic landscape of the human immune response, but a multi-omics view of immunity is necessary as mRNA profiles are not enough to understand immune activation^21, 22^. Transcript information can be leveraged to study the proteome, including identification of novel proteoforms resulting from alternative splicing. Proteoforms discovered by proteogenomics methodologies have already been found to have a role in immunological processes, for instance in immune-regulating micropeptides^23^ and tumor neoantigen production^24^.

Here, we stimulated peripheral blood mononuclear cells (PBMCs) with multiple microbial stimuli *in vitro/ex vivo* and performed long- and short-read RNA sequencing and secretome proteomics to gain insight into potential differences in immune response. We aim to provide insight into the immune transcriptome and proteome of immune cells during innate immune responses against a variety of pathogens.

## Material & methods

### Ex vivo PBMC experiments

Venous blood was drawn from five healthy donors^5^ and collected in 10mL EDTA tubes. Isolation of peripheral blood mononuclear cells (PBMCs) was conducted as described elsewhere^25^. In brief, PBMCs were obtained from blood by differential density centrifugation over Ficoll gradient (Cytiva, Ficoll-Paque Plus, Sigma-Aldrich) after 1:1 dilution in PBS. Cells were washed twice in saline and re-suspended in serum-free cell culture medium (Roswell Park Memorial Institute (RPMI) 1640, Gibco) supplemented with 50 mg/mL gentamicin, 2 mM L-glutamine and 1 mM pyruvate. Cells were counted using a particle counter (Beckmann Coulter, Woerden, The Netherlands) after which the concentration was adjusted to 5 × 10^6^/mL. *Ex vivo* PBMC stimulations were performed with 5×10^5^ cells/well in round-bottom 96-well plates (Greiner Bio-One, Kremsmünster, Austria) for 24 hours at 37°C and 5% carbon dioxide. Cells were treated with lipopolysaccharide (*E. coli* LPS, 10 ng/mL), *Staphylococcus aureus* (ATCC25923 heat-killed, 1×10^6^/mL), TLR3 ligand Poly I:C (10 µg/mL), *Candida albicans* yeast (UC820 heat-killed, 1×10^6^/mL), or left untreated in regular RPMI medium as normal control. After the incubation period of 24h and centrifugation, supernatants were collected and stored at -80°C until further processing. For the RNA isolation, cells were stored in 350 µL RNeasy Lysis Buffer (Qiagen, Rneasy Mini Kit, Cat nr. 74104) at −80°C until further processing.

### RNA and protein isolation

RNA was isolated from the samples using the RNeasy RNA isolation kit (Qiagen) according to the protocol supplied by the manufacturer. The RNA integrity of the isolated RNA was examined using the TapeStation HS D1000 (Agilent), and was found to be ≥7.5 for all samples. Accurate determination of the RNA concentration was performed using the Qubit (ThermoFisher).

We extracted the secretome of the 24-hour stimulated PBMCs. To 250 µL of supernatant, 250 µL buffer containing 10% sodium dodecyl sulfate (SDS) and 100 mM triethylammonium bicarbonate (TEAB), pH 8.5 was added. Proteins were reduced by addition of 5 mM dithiothreitol and incubation for 30 minutes at 55°C and then alkylated by addition of 10 mM iodoacetamide and incubation for 15 minutes at RT in the dark. Phosphoric acid was added to a final concentration of 1.2% and subsequently samples were diluted 7-fold with binding buffer containing 90% methanol in 100 mM TEAB, pH 7.55. The samples were loaded on a 96-well S-Trap^TM^ plate (Protifi) in parts of 400 µL, placed on top of a deepwell plate, and centrifuged for 2 min at 1,500 x g at RT. After protein binding, the S-trap^TM^ plate was washed three times by adding 200 µl binding buffer and centrifugation for 2 min at 1,500 x g at RT. A new deepwell receiver plate was placed below the 96-well S-Trap^TM^ plate and 125 µL 50 mM TEAB containing 1 µg of trypsin was added for digestion overnight at 37°C. Using centrifugation for 2 min at 1,500 x g, peptides were eluted in three times, first with 80 µL 50 mM TEAB, then with 80 µL 0.2% formic acid (FA) in water and finally with 80 µL 0.2% FA in water/acetonitrile (can) (50/50, v/v). Eluted peptides were dried completely by vacuum centrifugation.

### Long-read library preparation and sequencing

Libraries were generated from one donor using the Iso-Seq-Express-Template-Preparation protocol according to the manufacturer’s recommendations (PacBio, Menlo Parc, CA, USA). We followed the recommendation for 2-2.5kb libraries, using the 2.0 binding kit, on-plate loading concentrations of final IsoSeq libraries was 90pM (*C. albicans*, *S. aureus*, Poly(I:C), RPMI) and 100pM (LPS) respectively. We used a 30h movie time for sequencing.

The five samples were analyzed using the isoseq3 v3.4.0 pipeline. Each sample underwent the same analysis procedure. First CCS1 v6.3.0 was run with min accuracy set to 0.9. IsoSeq lima v2.5.0 was run in IsoSeq mode as recommended. IsoSeq refine was run with ‘--require-polya’. The output of IsoSeq refine was used as input for IsoQuant v3.1.2^26^ with GRCh38.p13 v43 primary assembly from GENCODE. The settings were set for full length PacBio data, and quantification included ambiguous reads. In IsoQuant, transcripts were considered novel if their intron chains did not match intron chains found in GENCODE annotation version 39.

Transcripts with fewer than 5 reads across all samples were excluded from further analyses (Supplemental table 1).

We sought to validate the novel transcripts identified using long-read sequencing using FANTOM5 CAGE data of CD14 monocytes (https://fantom.gsc.riken.jp/5/datafiles/latest/basic/human.primary_cell.CAGEScan/CD14% 2b%20monocyte%20derived%20endothelial%20progenitor%20cells%2c%20donor1.NCig100 41.11229-116C5.hg19.GCTATA.clusters.bed.gz) that allows for the identification of transcripts with a matching TSS from this 5’ sequencing data. Transcripts with novel 5’ were considered to be supported with a CAGE peak if within 150 basepairs from the TSS.

### Short-read library preparation and sequencing

RNA input was normalized to 200 ng for all samples/donors and libraries were generated using the QuantSeq 3’ mRNA-Seq Library Prep Kit-FWD from Lexogen (Lexogen) in accordance with the manufacturers’ protocol. In order to ensure high quality libraries, two separate preparations were performed, limiting the number of samples to 30 per preparation. End-point PCR was performed with 19 – 22 cycles, as indicated by a quantitative PCR on a 1:10 aliquot of a subset of double stranded cDNA libraries. Accurate quantification and quality assessment of the generated libraries was performed using Qubit dsDNA High Sensitivity assay (Thermo Fisher Scientific) and Agilent 2200 TapeStation (High Sensitivity D1000 ScreenTape, Agilent). Molarity of individual libraries was calculated using the cDNA concentration (Qubit) and average fragment size (TapeStation). Safeguarding sufficient read-depth for each sample, libraries were split in two separate runs. In each run, the baseline RPMI condition across all donors and time-points was included, in turn allowing sequencing bias assessment. The cDNA libraries of 35 samples were pooled equimolarly to 100 fmol. After a final dilution of both pools to a concentration of 4 nM, they were sequenced on a NextSeq 500 instrument (Illumina) with a final loading concentration of 1.4 pM.

FastQC v0.11.5 (Babraham Bioinformatics) was used to assess the quality of the obtained sequencing data, followed by removal of adapter sequences and poly(A) tails by Trim Galore! V.0.4.4_dev (Babraham Bioinformatics) and Cutadapt v1.18^27^. Since QuantSeq reads only provide coverage of the 3’ end of transcripts, we generated a set of transcripts representative of the full transcriptome by grouping transcripts based on unique 3’ sequences. Therefore, we separately mapped the filtered and trimmed reads to the long read transcriptome with Salmon v1.9.0 in mapping-based mode with decoys^28^.

### Differential expression analyses

To measure differential gene expression from long-read RNA sequencing, low abundance genes were filtered using a 10 CPM threshold with the conform package in python. Differentially expressed genes (DEGs) and transcripts were calculated for each condition versus control using the NOISeq R package^29^ from the abundances generated with isoquant. TMM normalization was chosen and q-value threshold for DE was set at 0.95.

DEGs were generated from the salmon-mapped short-read RNA sequencing data using the samples from the same donor using NOISeq^29^. The two control samples (RPMI) per donor were treated as technical replicates. TMM normalization was chosen and q-value threshold for DE was set at 0.95. We validated the DEGs detected from long-read sequencing with those generated with the short-read data by comparing the linear correlation of the log2fold change values for each condition combination between both datasets using the lm() R function.

The up- and downregulated DEGs per condition-control pair were analyzed for pathway enrichment separately using gProfiler^29^. We used Gene Ontology biological process and molecular function and TRANSFAC transcription factor motifs gene sets^30, 31^. A term size filter of between 100-500 was used to generate the final enrichment profiles.

### Isoform switching

A first-pass isoform switching analysis was performed using swanvis v2.0^32^. For a second-pass isoform switching analysis, the resulting gene-level isoform switch p-values were imported into IsoformSwitchAnalyzeR v1.16.0 package in R^33^. Thresholds for isoform switching were set at 10 DPI (differential percent isoform use) and nominal p-value <0.05. Sequences corresponding to the significant isoform switches were analyzed with CPAT v1.2.4^34^, hmmscan v3.3.2 with Pfam^35^, and SignalP5^36^ as a part of the IsoformSwitchAnalyzeR package.

Pathway analysis and gene network analysis of genes that were found to undergo isoform switching was performed in Cytoscape^37^. Default pathway analysis was performed, filtering for Gene Ontology Biological Process gene sets. An Enrichment Map was built from the enriched gene sets with a Jaccard similarity cutoff of 0.4^38^.

Genes found to undergo intron retention gains/losses and genes with domain gains/losses were separately analyzed using gProfiler. We used Gene Ontology Biological Process gene sets with a with a term size filter between 100-500 genes. We separately analyzed genes with domain gains or losses were using dcGOR^39^. We used the gene ontology molecular function gene sets with a term size filter between 100-500 genes.

### LC-MS/MS analysis

Peptides were re-dissolved in 20 µL loading solvent A (0.1% trifluoroacetic acid in water/acetonitrile) (98:2, v/v)) of which 4 µL was injected for LC-MS/MS analysis on an Ultimate 3000 RSLCnano system in-line connected to a Q Exactive HF mass spectrometer (Thermo). Trapping was performed at 10 μL/min for 4 min in loading solvent A on a 20 mm trapping column (made in-house, 100 μm internal diameter (I.D.), 5 μm beads, C18 Reprosil-HD, Dr. Maisch, Germany). The peptides were separated on a 250 mm Waters nanoEase M/Z HSS T3 Column, 100Å, 1.8 µm, 75 µm inner diameter (Waters Corporation) kept at a constant temperature of 45°C. Peptides were eluted by a non-linear gradient starting at 1% MS solvent B reaching 33% MS solvent B (0.1% formic acid (FA) in water/acetonitrile (2:8, v/v)) in 100 min, 55% MS solvent B (0.1% FA in water/acetonitrile (2:8, v/v)) in 135 min, 97% MS solvent B in 145 minutes followed by a 5-minute wash at 97% MS solvent B and re-equilibration with MS solvent A (0.1% FA in water).

The mass spectrometer was operated in data-dependent acquisition mode, automatically switching between MS and MS/MS acquisition for the 16 most abundant ion peaks per MS spectrum. Full-scan MS spectra (375-1500 m/z) were acquired at a resolution of 60,000 in the Orbitrap analyzer after accumulation to a target value of 3,000,000. The 16 most intense ions above a threshold value of 15,000 were isolated with a width of 1.5 m/z for fragmentation at a normalized collision energy of 28% after filling the trap at a target value of 100,000 for maximum 80 ms. MS/MS spectra (200-2000 m/z) were acquired at a resolution of 15,000 in the Orbitrap analyzer.

### Protein identification and quantification

Two search databases were constructed; one database for proteoform detection and one database for quantification. The database used for sensitive detection of proteoforms was generated using a slightly adapted version of the Long Read Proteogenomics pipeline by Miller *et al*^40^. Since the pipeline uses a different long-read transcriptomics tool, small syntax adjustments were made to accommodate the use of Isoquant output. Additionally, a custom script was written to have Isoquant output mimic the required input format. The pipeline generated a GENCODE-PacBio hybrid database. The proteome from *C. albicans* (taxon ID 5476) and *S. aureus* (taxon ID 1280) were downloaded from UniProt and added to the search database. The search database used for quantification was created by downloading the proteome from *H. sapiens* (taxon ID 9609), *C. albicans* (taxon ID 5476) and *S. aureus* (taxon ID 1280) from UniProt. Metamorpheus default contaminants were added to both search databases.

Mass spectra were identified using Metamorpheus v1.0.0^41^. The Human Proteome Project Mass Spectrometry Data Interpretation Guidelines version 3.0 were applied^42^. Quantification was performed using FlashLFQ v 1.2.4.294^43^ with all five individuals set as biological replicates and the two control (RPMI) samples per individual set as technical replicates. The following options enabled: normalization, shared peptide quantification, Bayesian fold change analysis, and match between runs (Supplemental table 2). An adapted version of SQANTI protein was used to search for novel peptides in the Metamorpheus identifications. Enrichment of secreted proteins was determined using the predicted secreted proteins from Human protein atlas^44^ as reference.

### Protein clustering

FlashLFQ raw protein expression values originating from the quantification database search were first square root transformed. To normalize for donor effects, the mean protein expression value per gene/individual was subtracted from all the expression values from the same gene/individual. Then z-score normalization was performed across all individuals per gene. K-means clustering was then performed using the kmeans() function in R with seed #82 and default parameters. We found four clusters to optimally represent the data according to the elbow plots (Supplemental figure 1). A heatmap was constructed with those clusters using the ComplexHeatmap package^45^. The proteins identifiers assigned to cluster #4 were converted to gene names and analyzed using gProfiler for enrichment analysis using both Gene Ontology Biological Process and Molecular Function gene sets. We further analyzed the protein found to form cluster 4 through a protein network analysis in Cytoscape^37^.

## Results

We stimulated PBMCs from five donors with four different microbial stimuli, mimicking bacterial (*E. coli* LPS, *S. aureus*), viral (Poly(I:C)) and fungal (*C. albicans*) infections. PBMCs were stimulated for 24 hours. RPMI incubation was used as a negative control (Figure 1A). To characterize full-length transcript structures, we performed long-read sequencing on PBMCs from one donor (Figure 1B). Additionally, shotgun proteomics data was generated from supernatants of the samples from all five donors. The proteomics data serves to corroborate differential gene/transcript expression and provide evidence of the protein-coding potential of novel transcripts identified through long-read RNA sequencing (Figure 1C). Short-read 3’ sequencing data of all five donors was generated to validate differential gene expression data generated from long-read RNA sequencing (Figure 1D).

**Figure 1:**
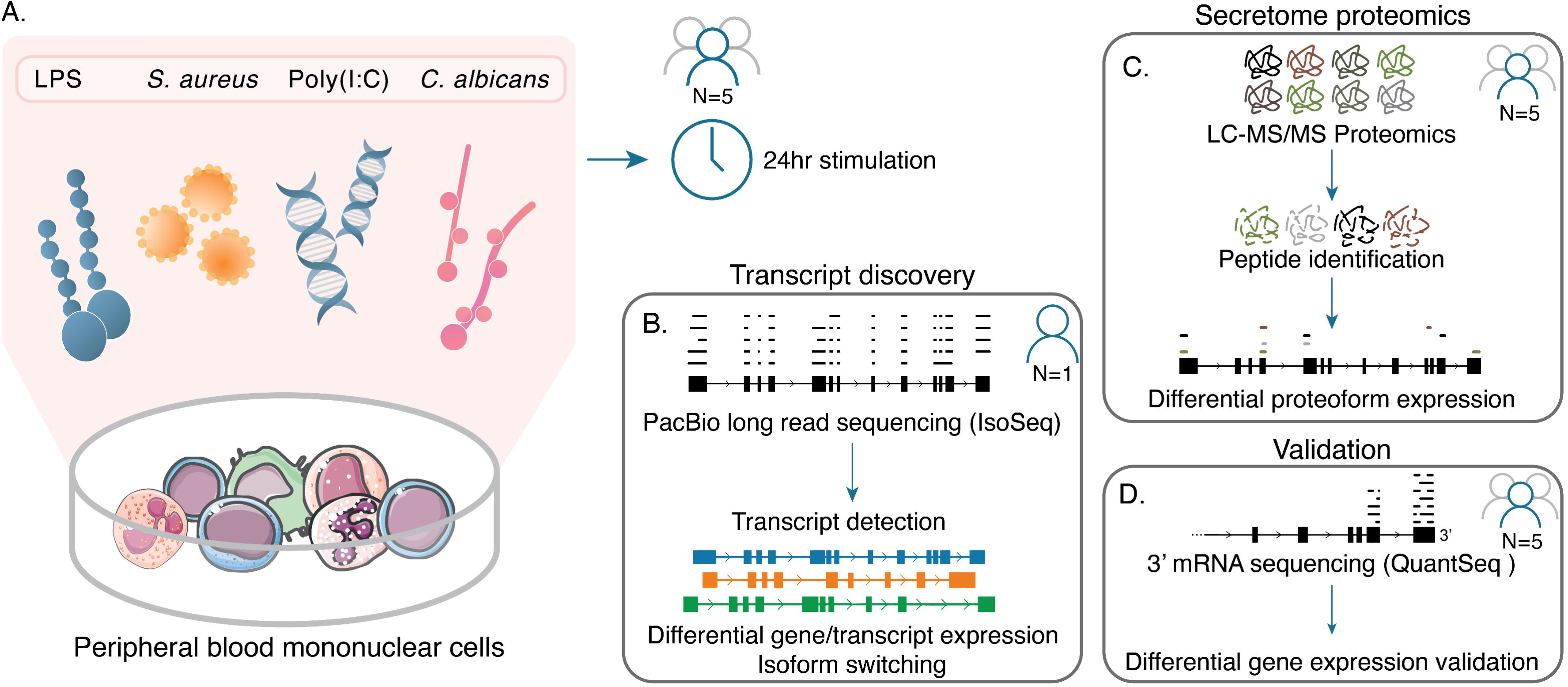
Experimental setup. A) Human peripheral blood mononuclear cells (PBMCs) isolated from five donors were exposed to four different pathogenic stimuli and analyzed after 24 hours. B) PacBio long read RNA-sequencing was performed on samples from one of the five donors. Long-reads were used to estimate differential transcript expression and isoform switching. C) The supernatant from all samples (all donors) was collected and peptides were detected to quantify protein levels in the secretome. D) Short read RNA sequencing (QuantSeq) was performed on all samples (all donors) and differential expression estimates were compared to those measured in long-read sequencing.

### Long read transcriptomes of both control and pathogen-stimulated conditions show novelty

Sequences detected using long-read sequencing were categorized in terms of novelty according to their intron chains. Transcripts are divided into three categories that encompass reference transcripts (GENCODE), novel in catalog (transcripts that contain annotated introns) and transcripts that are novel not in catalog (containing unannotated introns) (Figure 2A). We identified a total of 37,312 unique transcript sequences from 11,872 genes across all samples. The majority of transcripts were in protein coding genes (Supplemental figure 2A) including ∼10% immune-related genes (Supplemental figure 2B). We found 47.4% of detected transcripts to be novel, while these accounted for only 20.3% of the total reads (Figure 2B). The distribution of reads per novel transcript was similar to that of known transcripts with a slight skew towards lower abundance (Supplemental figure 3A). Exon elongations were the most observed feature distinguishing novel from known transcripts, occurring in nearly a third of the novel transcripts found in RPMI. This was similar for the stimulated conditions (Figure 2C, Supplemental figure 3B). The percentage of novel transcripts and transcript deviations were similar for all conditions (Figure 2D). To corroborate the existence of novel transcripts, we analyzed FANTOM5 CAGE peaks in the vicinity of the transcription start sites for novel transcripts with novel 5’ ends. We found 8,233 (51.3%) novel 5’ end transcripts across all conditions to be supported by a CAGE peaks from unstimulated human monocytes (within 150 nucleotides)^46^.

**Figure 2:**
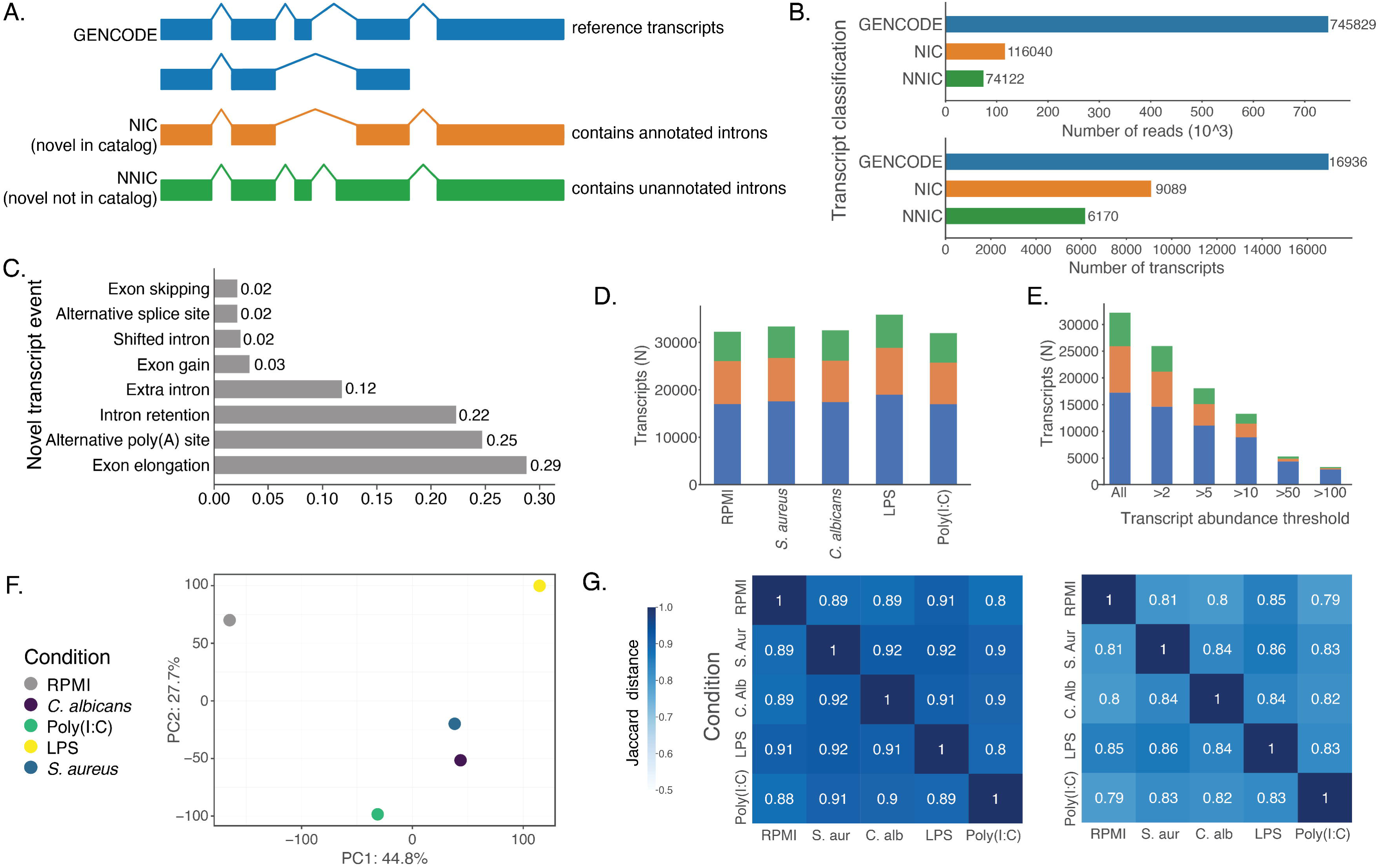
Transcriptome novelty in the control condition and comparison between stimuli transcriptomes in the five long-read samples. A) Transcript novelty categories. GENCODE (blue) is the set of all known reference transcripts. Novel in catalog (orange) contains a novel combination of annotated introns. Novel not in catalog (green) contains one or more unannotated introns. B) Reads (top) and unique transcripts (bottom) of events in each pre-defined transcript novelty category in RPMI. C) Novelty-inducing events occurring in the RPMI transcriptome. D) Unique transcripts by novelty category for each of the stimulus conditions. E) Unique transcripts by novelty category that remain at various transcript abundance thresholds in the C*. albicans* condition. F) Transcriptomes of the samples plotted on the first two principal components of PCA. G) Jaccard distances of genes (left) and known transcripts (right) of the transcriptomes, not considering transcript abundance.

Principal component analysis of the expression levels for each transcript indicated that stimulated conditions were more similar to each other than to RPMI. *S. aureus* and *C. albicans* were most similar to each other (Figure 2F). Genes and transcripts expressed were similar in the stimulated conditions with average Jaccard similarity indices of 0.9 and 0.82 for genes and transcripts, respectively (Figure 2G). Novel transcripts had similar Jaccard indices to each other than for known transcripts (not shown). Differential expression analysis yielded an average of 949 DEGs and 2,076 differentially expressed transcripts per condition (Supplemental figure 4, Supplemental table 3-4).

We validated the DEGs through 3’ transcript counting (QuantSeq)^47^. We gathered a set of representative transcripts based on sequence differences at the 3’ end of transcripts (29,760 transcripts, 79.8% of total) and investigated the correlation of differential expression in the long-read sequencing data with the separately generated short read dataset of the same donor. The DEGs that overlapped between both datasets correlate well (R^2^ 0.62-0.81). Best matching pairs of stimulated conditions between the short- and long-read confirmed the concordance of both sequencing approaches (Supplemental figure 5, Supplemental table 5).

### Pathogen stimuli display upregulation of different pathways

Differential gene expression analysis using the long-read sequencing data resulted in a total of 1,733 genes that were differentially expressed in stimulated conditions compared to control. We performed pathway analysis for each condition using gProfiler^48^ (Supplemental table 6). By overlapping the gene sets enriched in each of the four conditions, we discerned biological processes/functions specific to certain pathogen-stimulated conditions. There are a lot of constants in host response regardless of the pathogen, and indeed the largest set of pathways was in the overlap between all stimulated conditions (211 pathways, Figure 3A). This set has an enrichment of genes involved in type II interferon (IFN-γ) responses. Genes involved in tertiary and specific granules, which play a role in the defense against pathogens were found to be enriched among upregulated genes in all conditions. Surprisingly, we also find these and related gene sets to be enriched among downregulated genes as a result of *S. aureus* and Poly(I:C) stimulation, potentially a result of the regulation of the inflammatory response. Further gene sets included the response to molecules of bacterial origin (including LPS), innate immune response signaling such as PRR signaling, antigen processing and presentation and IL-1 production (Figure 3B).

**Figure 3:**
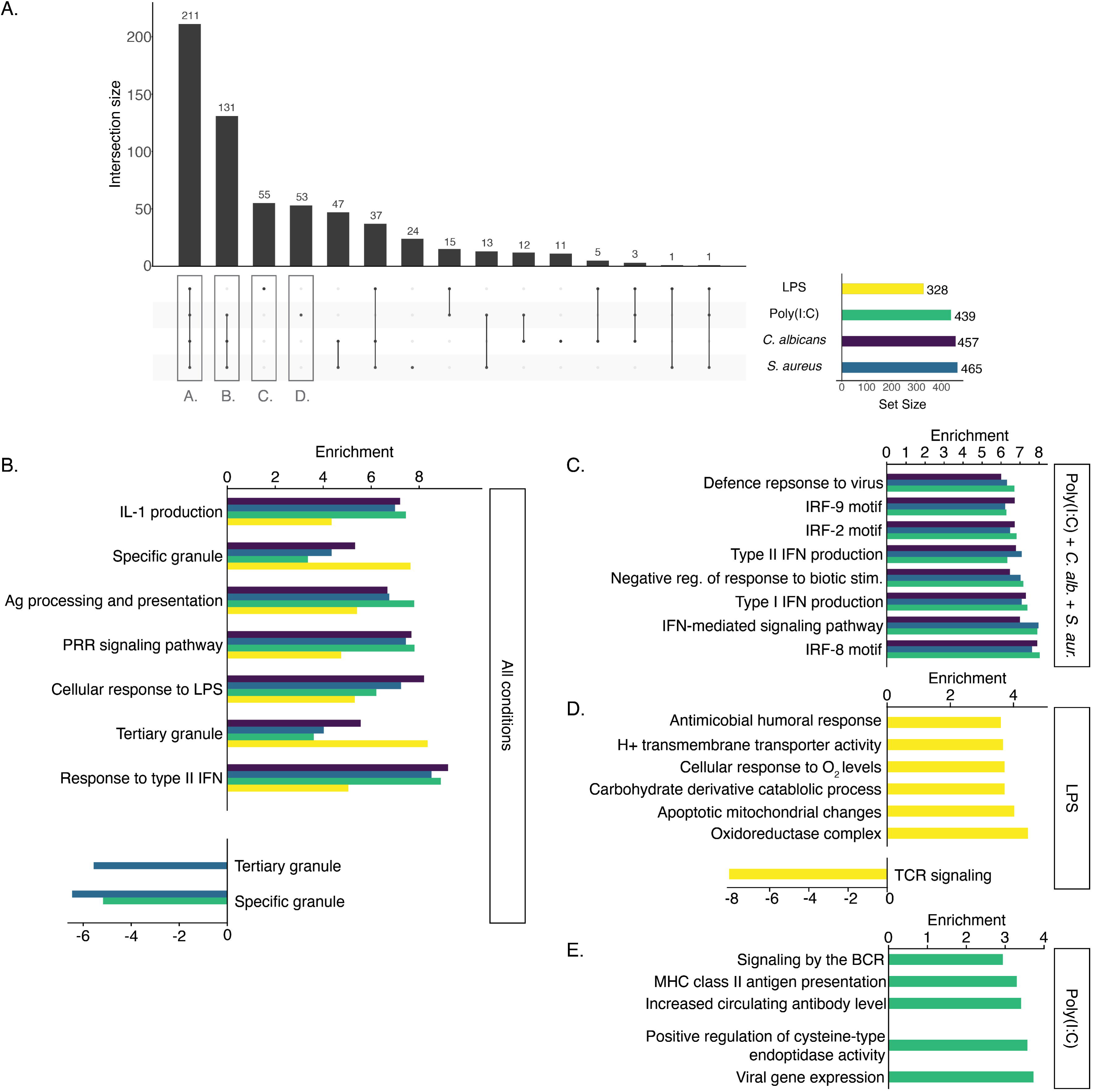
Differential pathway analysis originating from differentially expressed genes on the RNA level. A) Overlap between enriched pathways generated from the differentially expressed genes from the four conditions. B) Selected pathways found to be enriched for all conditions, C) three of the four conditions (Poly(I:C), *C. albicans* and *S. aureus*), D) specifically for LPS and E) specifically for Poly(I:C).

Some pathogen-stimulated conditions had more enriched pathways in common than others. There was a notable overlap of 131 gene sets enriched in *C. albicans-, S. aureus-* and Poly(I:C)-stimulated conditions. Some of these were common to the set overlapping between all conditions, such as interferon responses. The LPS-excluding set showed particular enrichment related to viral processes such as the defense against viruses, regulation of the viral lifecycle, likely due to interferon-stimulated gene expression, such as *STAT1, OAS1/3, OASL and IFIH1*. Also, transcription factor binding matches (TRANSFAC) such as *IRF-2, 5, 8* and *9* were enriched, reflecting downstream signaling through various signaling pathways leading to the regulation of the production of interferons and immune cell development (Figure 3C)^49^.

LPS and Poly(I:C) were the 2 stimuli with the most enriched pathways unique to a single stimulus. For 55 gene sets unique to LPS, there was a downregulation of T cell receptor signaling, in part due to the downregulation of *CD4* expression, which has previously been described as a result of endogenous production of TNF-α and IL-1β as a result of LPS stimulation^50^. We further found an upregulation of gene sets involved in metabolic processes such as oxidoreductase complexes and cellular responses to oxygen, possibly reflecting metabolic changes previously described to occur in immune cells such as monocytes upon LPS stimulation^51^. Furthermore, there was an upregulation of genes involved in humoral immune responses (Figure 3D). For 53 gene sets enriched uniquely in Poly(I:C), we found functions including viral gene expression, apoptosis related signaling (regulation of cysteine-type endopeptidase activity) and B-cell related gene sets such as increased antibody levels and BCR signaling. Finally, there was an enrichment of MHC class II antigen presentation (Figure 3E).

### Isoform switches highlight transcriptome differences between conditions and control

Isoform switching (IS) genes are defined by a change (increase/decrease) of expression of a particular transcript isoform as measured by percent of total reads for a gene. In different samples/conditions, a particular transcript isoform may comprise a different isoform fraction (dIF) value for a given gene. Here, a change of at least 10% (0.10 dIF) in control and the opposite change (decrease/increase) of expression of a different transcript isoform in the same gene of at least 10% in the pathogen-stimulated condition is considered an IS.

A total of 999 IS were detected in 398 genes. Nearly half (N=192, 48.2%) of these IS genes were unique to their respective stimulus conditions, while 10.3% were found in all conditions (N=41) (Figure 4A, Supplemental table 7-8). The majority of genes demonstrating IS were not differentially expressed in their respective conditions (327 genes; 77%). Most genes that were found to undergo IS displayed only one IS instance (Supplemental figure 6A). Pathway analysis of genes undergoing IS were enriched for gene sets involved in metabolic processes, mRNA splicing, protein transport and catabolism. Furthermore, immune and stress-related pathways such as MHC type I antigen processing and transport through vesicles, inflammasomes, oxidative stress and apoptosis were found to be represented in genes undergoing IS (Figure 4B, Supplemental tables 9-13).

**Figure 4:**
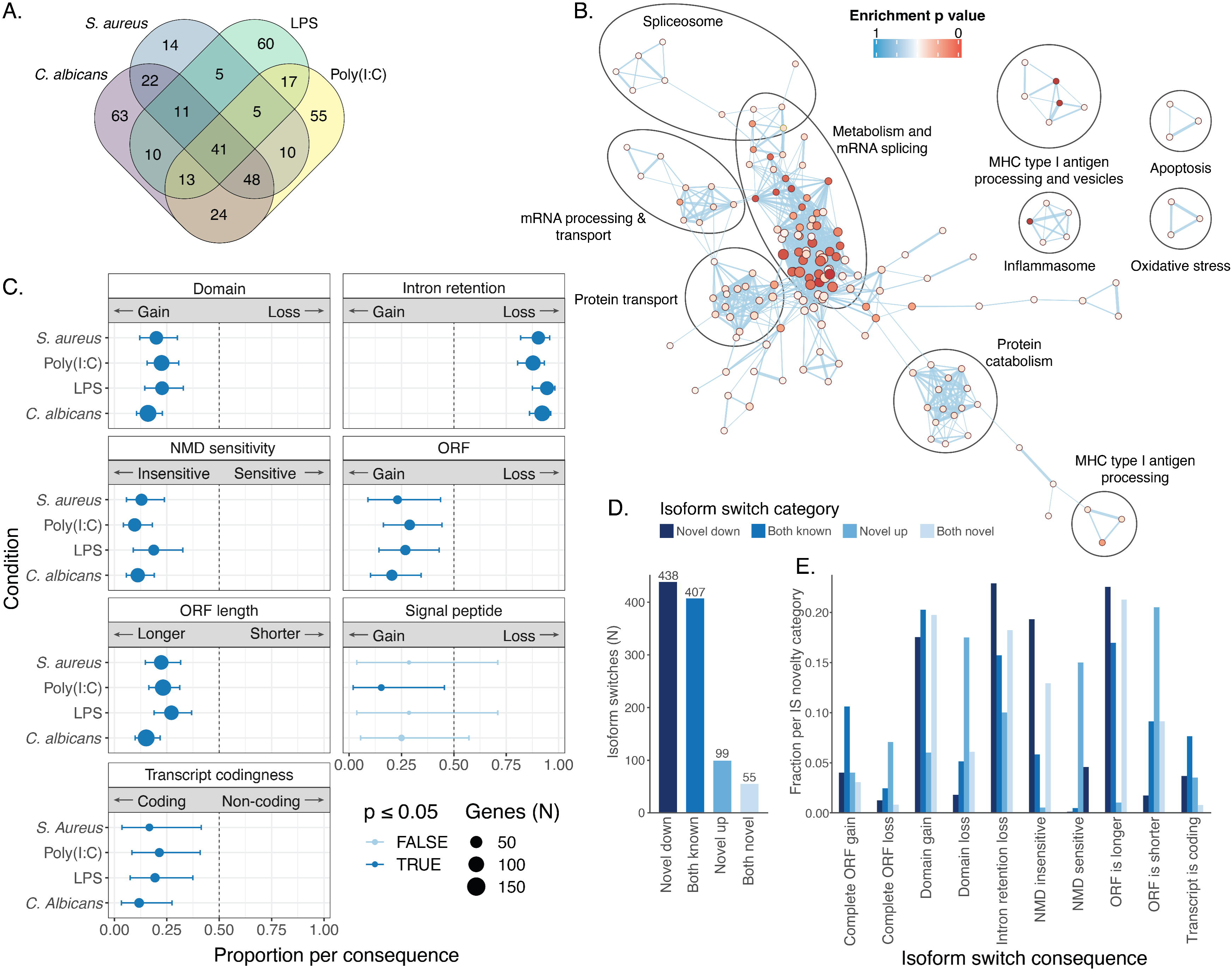
Isoform switching induced by pathogen stimulation. A) Overlap of isoform switching genes between the four stimulus conditions. B) Pathway network analysis derived from genes found to undergo isoform switching (IS) upon pathogen stimulation. Each pathway is colored by p value, where a darker red indicates a lower p value. C) Proportions of total IS events in each stimulated condition per IS consequence. D) Number of IS by category of switch pairs. Categories are defined by involvement of novel transcripts in a given IS. “Novel down” indicates that the isoform switched from a higher proportion of the novel transcript in control to a higher proportion of a known transcript in the stimulus condition. “Both known” indicates that the IS occurs between 2 reference transcripts. E) Fraction of each transcript novelty combination per IS consequence. Normalized by total number of IS events per novelty category.

We sought to understand the molecular consequences of IS upon pathogen stimulation by categorizing the differing features of the isoform pairs involved in the switch. Each of the IS was annotated with one or more of the following predicted protein characteristics: change in ORF length, ORF gain/loss, domain gain/loss, NMD sensitivity, intron retention (IR) gain/loss, coding probability (ORF presence), and signal peptide gain/loss. These consequences are not independent and often multiple consequences could be attributed to one IS (Supplemental Figure 6B). We observed general IS trends on a genome-wide scale (Supplemental figure 6C, Supplemental table 14). Strikingly, we found IR loss to be the most common consequence of IS in this dataset. Isoforms with retained introns comprised a higher isoform fraction for genes in the control condition, while their respective intron-excluding counterparts had a higher isoform fraction for genes in the pathogen-stimulated conditions. Genes displaying loss of IR were enriched for pathways involved in mRNA processing, including spliceosome-related gene sets, antigen processing and IL-1 production (Supplemental figure 7). IR has previously been described as a regulatory mechanism of RNA processing, splicing, vesicle transport and type I interferon production in the development of various immune cell types, including macrophages^52, 53^, granulocytes^54^ and B cells^55, 56^. Our findings support previously described associations of IR losses in immune-related processes, and adds new genes regulated by IR loss during immune responses (Supplemental figure 7, Supplemental table 15).

In addition, we found a higher proportion of transcripts to have domain gains than domain losses. This could indicate that stimulation by a pathogen causes a gene to switch expression to a transcript isoform that codes for a protein with an extra function. Other observed trends included longer ORFs and NMD insensitivity in transcript isoforms induced by pathogen stimulation (Figure 4C).

Since the addition or loss of domains could directly reveal protein function changes, we explored the IS that had this consequence type. We found that genes with domain gain/loss (N=158, Supplemental table 16) were enriched for involvement in various catabolic processes. We also found enrichment of T cell activation genes, an effect previously described as a functional consequence of CD8+ T cell co-stimulation^57^. Other enriched gene sets include leukocyte cell-cell adhesion and activation and general innate immune response genes (Supplemental table 17). When looking more specifically at the molecular functions of the gained domains themselves, we found an enrichment of domains with potassium channel regulator activity, kinase- and transferase activity concerning phosphorus-containing groups and nucleic acid binding. These results potentially indicate functional and cell-type specific effects of domain gains as a result of IS in immune responses (Supplemental figure 8, Supplemental table 18).

Novel transcripts play an important role in IS. Of the 999 IS, more than half (N=592) had at least one novel transcript involved in the IS. In most cases (N=438), the switch was from a novel transcript to a known transcript (Figure 4D, Supplemental table 19). Compared to IS cases where only known transcripts were involved, the IS consequences were more often NMD insensitivity and IR loss (Figure 4E). Conversely, shorter ORFs, domain losses and NMD sensitivity were more common effects when the IS was from a known to a novel transcript isoform. In conclusion, the unstimulated condition is characterized by the presence of many novel transcripts with retained introns, which are difficult to detect with short read sequencing. IR is likely a mechanism to prepare a cell for fast action after an immune stimulus, when splicing of the retained intron could quickly generate a functional transcript with coding potential, which has been for instance been described in CD4+ T cells^58^.

### A novel read-through transcript including *CARD16* and *CASP1*

As an example of a remarkable finding with possible biological impact once validated, we identified a read-through transcript that includes both *CARD16* and *CASP1* (Figure 5A). Read-through transcripts involve transcription that extends beyond the normal polyadenylation site (PAS), terminating at the PAS of an adjacent gene or other nearby locus^59^. These transcripts have been found to be expressed in specific circumstances, including malignancy and infection^59, 60^. This particular novel transcript encompasses the coding region of *CASP1* and has an extended 5’ UTR which spans *CARD16,* and thus contains two ORFs. This IS was annotated as an intron retention loss, as the novel transcript loses an intronic region in its 3’ UTR (Figure 5B). Both the known and novel transcripts in this IS are predicted to be coding (both 100%). *CASP1* was found to be differentially expressed upon Poly(I:C) stimulation (log2FC 1.73, p=0.049; Figure 5C). The isoform expression of the known transcript was found to decrease upon Poly(I:C) stimulation, while the novel transcript was found to increase (Figure 5D). This is further reflected in the isoform fraction, increasing from 8.3% to 24.8%, while the known transcript decreased from 85.5% to 74.2% (Figure 5E).

**Figure 5:**
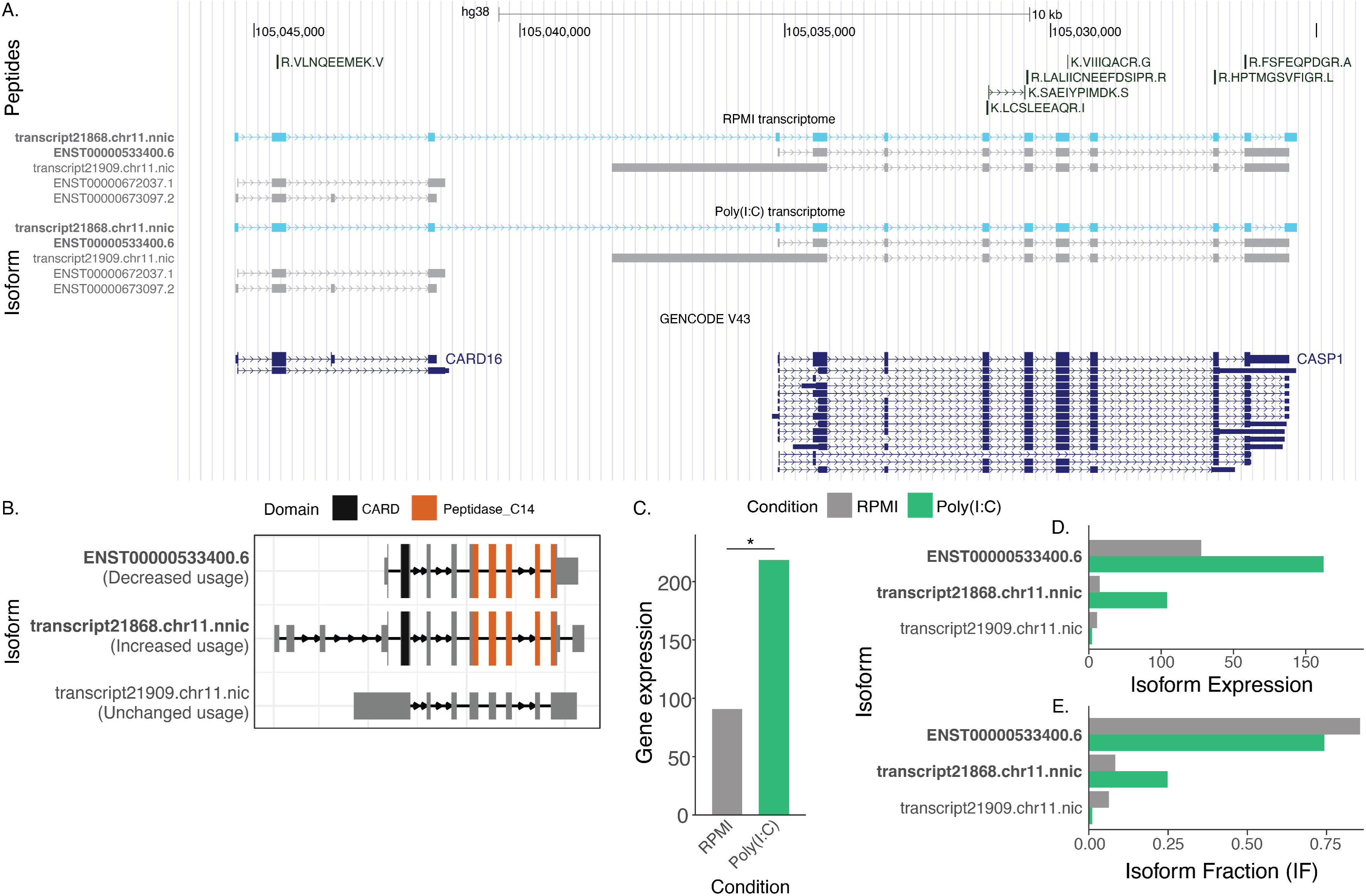
*A novel readthrough transcript of CASP1.* A) USCS genome browser track of the transcripts detected in the control condition (RPMI) and stimulated condition (Poly(I:C)). The novel readthrough transcript containing both CASP1 and CARD16 is presented in light blue. Known transcripts in in GENCODE are presented below. B) Representation of the domains in the novel *CASP1* transcript, indicating that *CARD16* is entirely included in the 5’ UTR of the transcript. C) Gene and transcript expression and isoform fraction of the *CASP1* transcripts that were detected.

*CARD16* and *CASP1* both have a function in proinflammatory IL-1β signaling, where *CARD16* has been shown to play a role in *CASP1* assembly, although there remains discussion on the exact regulatory effect of *CARD16* on this process ^61, 62^. We have identified an IS specifically for Poly(I:C) stimulation, where a novel transcript of *CASP1* was found to harbor *CARD16* in its 5’ UTR was upregulated upon stimulation. This finding could suggest a novel molecular mechanism in IL-1β signaling, potentially through the regulation of *CASP1* by its regulator *CARD16*.

### A novel coding transcript of *NFKB1*

We identified a novel *NFKB1* transcript that demonstrated IS in all four conditions. This novel transcript was shorter than the canonical transcripts (Figure 6A). Further analysis revealed that the novel transcript start site was supported by multiple nearby CAGE peaks (Figure 6B). Strikingly, this novel transcript lacks a part of its Rel homology domain, a conserved domain responsible for functions such as dimerization and DNA binding (Figure 6C)^63^. *NFKB1* was not found to be significantly differentially expressed, although gene expression was found to be higher in pathogen-stimulated condition compared to unstimulated condition (only *C. albicans* shown, Figure 6D). The expression of the novel transcript was found to increase upon pathogen stimulation (Figure 6E). This is reflected in the isoform fraction, which increases from 23.5% to 50.7%, while the known transcript decreases from 39.0% to 21.2% (Figure 5F).

**Figure 6:**
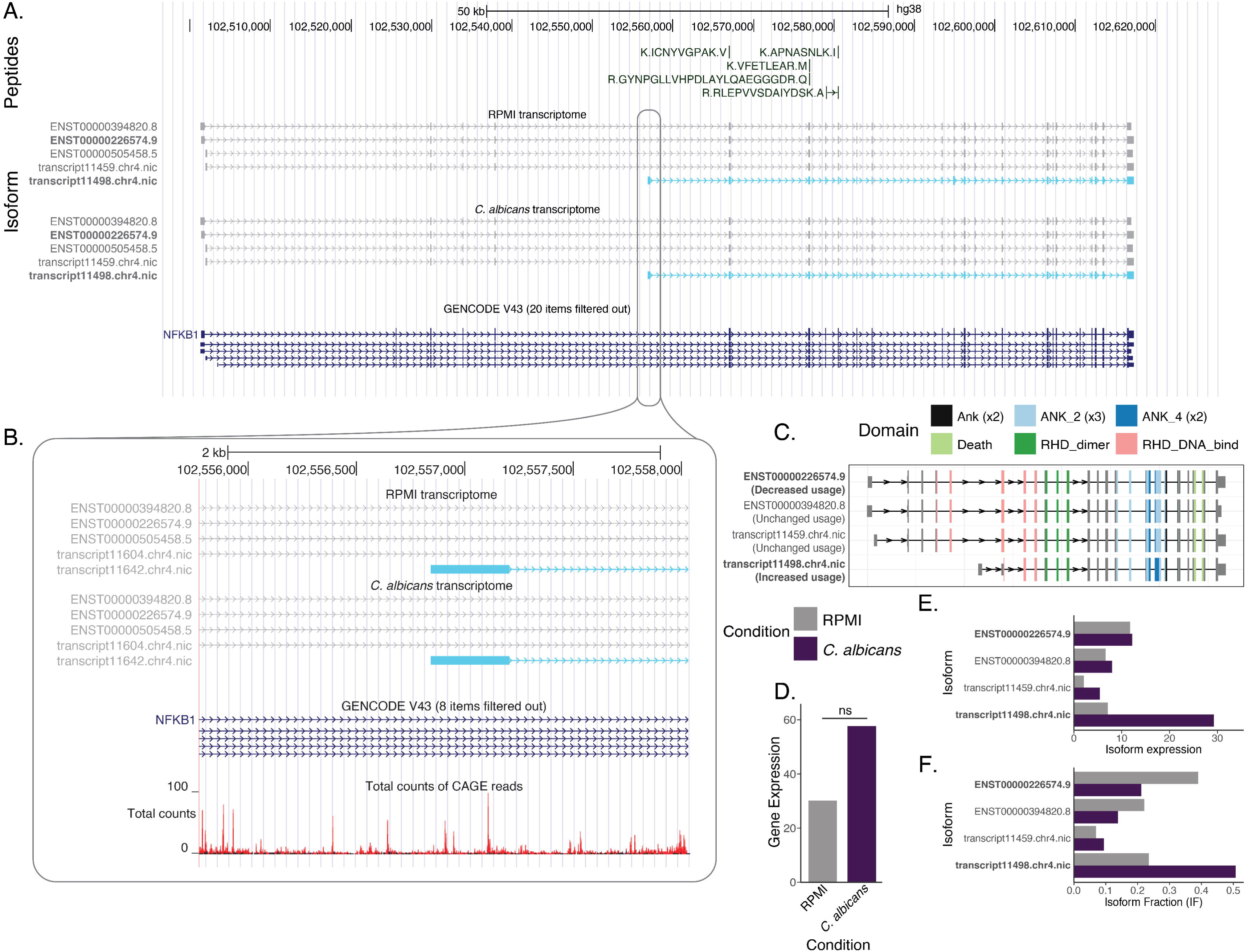
A novel transcript of *NFKB1.* A) UCSC genome browser track of the transcripts detected in the control condition (RPMI) and stimulated conditions. The novel transcript is presented in light blue. Known transcripts in GENCODE are presented below. B) Zoomed view of the transcription start site of the novel transcript with CAGE peaks (monocyte) in this region. C) Representation of the domains in the known and novel *NFKB1* transcripts that were detected. D) Gene and transcript expression and isoform fraction of the *NFKB1* transcripts that were detected.

*NFKB1* plays a central role in immune responses, regulating the response to infections through transcriptional activation^57^. Furthermore, the Rel homology domain region is known to harbor disease-causing variants responsible for common viable immunodeficiency (CVID)^64^, highlighting the importance of this domain in normal B cell function. This finding could suggest a novel regulatory mechanism of *NFKB1*.

### Isoform switching in *CLEC7A* and *OAS1* in a stimulus-specific manner

We sought to identify genes with stimulus-specific IS patterns. We identified an IS in *CLEC7A,* which codes for Dectin-1, a receptor that recognizes fungal glucans, triggering the immune response^65^. While this gene was not differentially expressed upon pathogen stimulation, we did identify an IS specific to *C. albicans* stimulation in this gene, involving a decrease in expression of an NMD sensitive transcript, with the increase in expression of the canonical coding transcript and a non-coding transcript (Supplemental figure 9A). In contrast, IR loss in *CLEC7A* was previously identified as a result of stimulation with multiple pathogen stimuli in monocytes. Additionally, no difference in gene expression levels was found between IAV-stimulated and resting cells, where the change in splicing was most pronounced^66^. While we find IR loss in *CLEC7A* specifically upon *C. albicans* stimulation, this could therefore also indicate a shared transcriptional response to pathogen stimuli.

Additionally, we identified an IS involving *OAS1*, which is involved in antiviral immunity. This gene is differentially expressed in response to *C. albicans, S. aureus* and Poly(I:C) (Supplemental figure 9B). We identified an IS in this gene resulting in an intron loss and a domain gain. This IS was only observed for Poly(I:C) stimulation, potentially indicating this transcript is necessary for antiviral immune responses (Supplemental figure 9C). Previous work has identified common *OAS1* haplotypes responsible for a decrease in protein abundance through the expression of NMD sensitive transcript p42, which contributes to COVID-19 severity^67^. We find this transcript to be downregulated upon Poly(I:C) stimulation. However, the transcript we find to be upregulated lacks the prenylation site needed for antiviral function, as shown for p46^68^.

### Detecting secreted peptides

We sought to obtain evidence of the protein-coding potential of novel transcripts found through long-read RNA sequencing. Mass spectrometry was performed for 30 secretome samples from five donors’ stimulated PBMCs, which includes the samples from the individual for which long read RNA sequencing was performed (see methods). These include 2 control samples and 1 of each 24-hour pathogen stimulation condition for each individual.

We designed a search database comprising all proteins that we suspected could be in the sample. This includes the GENCODE human proteome, the proteomes of the pathogens used, as well as ORFs derived from novel transcripts found using long-read RNA sequencing. Novel transcripts do not always correspond to novel ORFs; 32% of the novel transcripts had an ORF that was present in the GENCODE reference database (Supplemental Figure 10). In the collection of 30 samples, a total of 38,703 peptides from 15,964 proteins were identified. We found 404 (7.37%) of identified proteins were known to be secreted according to the human protein atlas, which constitutes a significant enrichment (OR=2.12, p=3.88x10^-21^, Fisher’s exact test). We did not detect microbial proteins in the samples. Many of the novel ORFs predicted from the transcriptome have high similarity to GENCODE ORFs, resulting in a small number of novel peptides that could uniquely identify these. After rigorous filtering, we were unable to confidently identify peptides that mapped uniquely to the predicted novel ORFs.

### Wider deviations in expression in the secretome

To assess whether differences in transcript expression resulted in differences in the amounts of secreted proteins, we performed a label-free quantification of the proteins in the cells’ supernatants. Using PCA, we found that a large portion of variation in the proteome was explained by inter-individual differences and that these differences were larger than the differences induced by the immune stimuli (Supplemental figure 11).

We found a total of 418 differentially expressed proteins (DEPs) between the stimuli and control when controlling for individual variation. Differential protein expression was not equally distributed between stimuli with over a third (N=131) of the DEGs unique to Poly(I:C) stimulation (Supplemental figure 12A). With the exception of the *S. aureus* condition, more proteins were significantly downregulated than significantly upregulated in the secretome (Supplemental figure 12, Supplemental table 20). We found few overlapping proteins per condition, which could indicate either a high specificity in response to different pathogens or a lack of protein secretion in a subset of samples.

To determine which explanation is more likely, we visualized the specific (groups of) proteins associated with response stemming from the stimuli. We clustered protein expression values normalized by individual and stimulus (Figure 7A, Supplemental table 21). The clustering revealed a separation between poly(I:C) samples and the rest of the stimuli. *C. albicans* showed a large overlap with poly(I:C) in the protein expression profiles. Some *C. albicans* samples were grouped with poly(I:C) samples, which confirms the results from the differential protein expression analysis (34 common DEPs, Supplemental figure 12A). Other stimulus conditions could not reliably be separated from RPMI.

**Figure 7:**
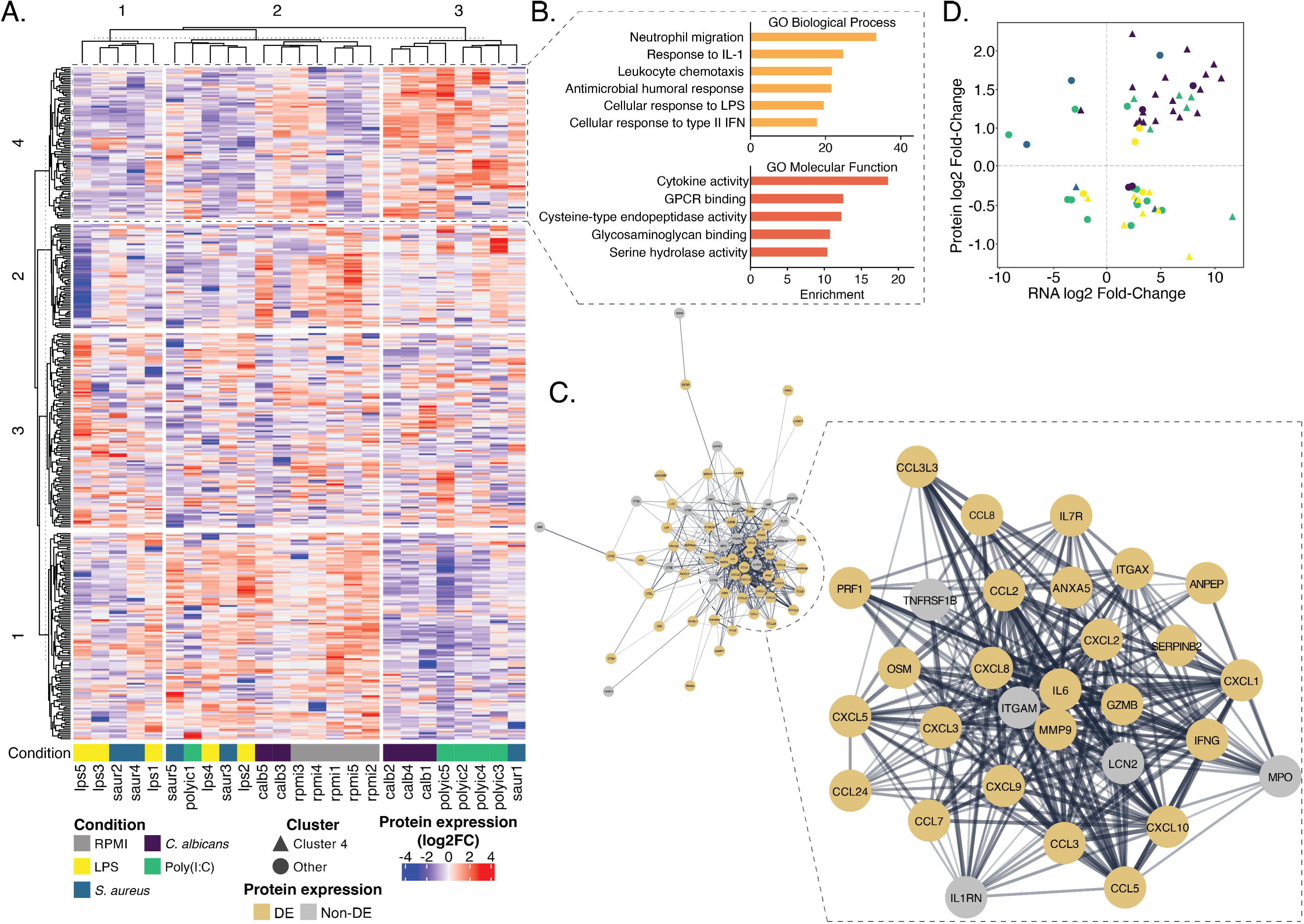
Protein expression in the secretome. A) Normalized protein expression detected from five donors in the five conditions. Donors are denoted with numbers 1 through 5. Clusters originating from kmeans clustering are shown in the heatmap. B) Gene ontology Biological process (top) and molecular function (bottom) pathways of proteins found in cluster 4. C) Clustering of genes found in cluster 4. Genes are found to be differentially expressed on the protein level are in color, others greyed. D) Fold change of gene differential expression versus fold change of protein DE for genes that were differentially expressed on both levels, colored by stimulus. Triangle-shaped points correspond with cluster 4 genes from A. Genes with concordant protein and RNA expression are in the upper right and lower left quadrants.

We identified a cluster of proteins that are highly expressed in Poly(I:C) and *C. albicans* (cluster 4, Figure 7A). This group of proteins is enriched for genes with functions in leukocyte migration and chemotaxis, exemplified by neutrophil migration. We identified further enrichments of gene sets involved in the response to IL-1, humoral antimicrobial response, and cellular responses to LPS and type II interferons. Analysis of the molecular functions of these genes indicated an enrichment of cytokine activity and receptor binding, GPCR receptor binding and various catalytic functions, likely due to immune cell differentiation and immune responses involving the degradation of extracellular matrix proteins during immune cell migration^69^ (Figure 7B, Supplemental table 22). We further assessed the proteins in cluster 4 through a gene network analysis (Figure 7C, Supplemental table 23-24). Of the 84 proteins in this network, 61 were differentially expressed on the protein level (72.6%, any condition). Of these DEPs, 18 are involved in cytokine signaling (29.5%), of which 13 genes are chemokines (71.2%). A high proportion of proteins are found in the extracellular region (n=47, 77.0%), for instance through secretion in granules. The biological functions of the DEPs in cluster 4 reflect those found for the complete set of proteins in cluster 4, mainly corresponding to pathways associated with functions in neutrophil migration and chemotaxis (Supplemental table 25). As these pathways are not necessarily specific to these two stimuli, this may indicate Poly(I:C) and *C. albicans* may be more effective at eliciting differential protein secretion or have less delay in secretion compared to the other stimuli.

### Comparison with RNA expression

As established earlier, a multi-omics approach is currently the best way to understand the human immune response. Correlation between the RNA and protein levels, or lack thereof, can provide important clues about the host response to pathogens. To assess the correlation of differential gene and protein expression levels, we assessed the concordance of differential expression on the RNA and protein level. This metric corresponds to the percentage of genes for which differential expression on both levels matched in directionality (out of all genes where DE was observed on both levels) (Figure 7D, Supplemental table 26).

We observed an overall poor concordance of directionality and fold change of expression levels at the RNA and protein levels in the different stimulus conditions, with the exception of *C. albicans* with 73% overall concordance. We overlaid the genes in group 4 from our clustering analysis with the genes found to be DE on both RNA and protein levels. There was an overrepresentation of the genes in this cluster in the total group of dual-level DE genes (OR=6.99, p=4.813e-16). Further analysis of concordant differential expression matches arising from proteins in cluster 4 (triangles in Figure 7D), we observed high concordance in the genes induced by *C. albicans* and/or Poly(I:C). Directionality concordance for Poly(I:C) and *C. albicans* for genes in in cluster 4 was significantly higher than overall directionality concordance (p=0.0313 Poly(I:C), p=0.0003 *C. albicans*, Fisher’s test one-tailed). The cluster 4 proteins in the LPS and *S. aureus* conditions are in the lower right quadrant, indicating that the increase of RNA translated into a decrease of secreted proteins for these genes (Figure 7D).

We hypothesized in the IS analysis that a major regulatory mechanism in the host response to pathogens was the loss of intron retention for rapid protein generation. We cross-referenced the secreted proteins to support this conjecture. By overlapping upregulated isoforms from intron retention loss events, we found 20 cases from 7 genes (Supplemental table 27). Of these genes, 2 were upregulated on the protein level, supporting our hypothesis. The genes were *GZMB* and *B2M*, which are important immune-regulatory genes that are both secreted^70, 71^. Considering the remaining 5 genes that were *downregulated* on the protein level, however, this is not convincing evidence that intron retention loss in general provides a rapid increase of protein production.

## Discussion

The identification of novel transcripts and subsequent production of additional protein isoforms could help identify molecular mechanisms that play a role in various biological processes, including immune responses. Various immune system processes have previously been found to be regulated by alternative splicing ^72, 73,74 75^. Immune responses display significant inter-individual differences. Donor-specific effects such as sex and ancestry have been shown to significantly influence the transcriptome. Previous studies have further shown the impact of QTLs in the heritability of cytokine production capacity^5, 76–78^. However, the effect of these processes on host defense mechanisms against pathogens, together with the large inter-individual differences in transcription and protein expression, remain to be elucidated.

We have generated a long-read transcriptome of pathogen-challenged primary immune cells (PBMCs) together with the secreted proteome to investigate mechanisms underlying immune responses during infection. We described the accurate identification of known and novel transcripts in both control and pathogen-challenged conditions. Of these transcripts, we identified a subset that is differentially expressed as a result of pathogen stimulation, which we validated by short read RNA sequencing data (including 4 additional individuals) and publicly available CAGE data from neutrophils.

We examined isoform switching that occurred as a result of pathogen stimulation, insight into transcripts that may play a role in pathogen responses. On a genome-wide level, widespread intron retention losses were observed. Retained introns that rendered the transcript unusable in the control condition were spliced out as a result of microbial stimulation; a trend we observed in all conditions regardless of microbe. We postulate that these are examples of unproductive splicing in unstimulated cells switching to productive splicing after stimulation enabling fast production of proteins relevant for the immune response. Genes that undergo intron retention loss mainly have functions in mRNA splicing and processing and in immunity. Tissue- or cell-type specific unproductive splicing has been widely observed as an autoregulatory process for mRNA splicing factors^79^, which is supported by our data in immune cells. We were however not able to confirm changes in protein expression of genes that underwent IR losses using our secretome proteomics data, likely because these proteins are not generally secreted. A couple of pertinent examples have been illustrated in greater detail. We identified an IS specific to the viral stimulus that involves a novel read-through transcript of *CASP1* and *CARD16*. We found an instance of IS to a novel *NFKB1* transcript with a shortened DNA binding domain that was found in all four conditions. Additionally, we describe IS in *CLEC7A* and *OAS1* for *C. albicans* and Poly(I:C), which highlight stimulus-specific alternative splicing. Taken together, these results highlight the potential for long-read sequencing to accurately resolve novel transcripts with potential relevance in immune responses, including intron retention loss events that are generally difficult to detect using short-read sequencing.

The extent to which conclusions can be drawn about immune response mechanisms is limited by the low sample size for long-read sequencing. In this explorative study meant to provide insights into the novel technical possibilities utilizing latest sequencing approaches, we generated long-read sequencing data for only a single individual because of the expensive nature of this technology in combination with the required sequencing depth and the number of conditions studied. This design did not allow us to investigate the inter-individual differences in the transcriptome. Novel transcripts that were detected could thus be specific to this individual. Future follow-up including the sequencing of more individuals using accurate long-read sequencing methods and functional studies could provide additional insight into the more general relevance of these transcripts in immune responses. This study focused on the appraisal of the transcriptome and proteome in PBMCs, which consist of multiple cell types. Use of freshly isolated PBMCs accurately represents the complete immune cell population in the peripheral blood and allows for communication between cell types during pathogen stimulation, thereby potentially giving an accurate representation of this cell population *in vivo*. However, no information on cell type specificity of transcripts is available. This could be resolved by recent developments in single cell long-read sequencing^80^.

The proteome, in contrast, was generated for all samples from all 5 individuals and highlighted significant differences between the secretome of individual donors, before and after response to immune stimuli. Concordance between the transcriptome and proteome levels was high in Poly(I:C) and *C. albicans*, and lower in LPS and *S. aureus*. We found that genes with high correlation on the RNA- and protein levels form a cluster of protein expression, separating the former two stimuli from the latter. These proteins are enriched for secreted immune-related proteins, indicating that pathogen stimulation successfully led to secretion of relevant proteins. This would indicate that cells have responded faster to the Poly(I:C) and *C. albicans* stimuli than to the LPS and *S. aureus* stimuli, because RNA and protein were isolated simultaneously from our samples. Delay in protein production after expression of an mRNA may partially explain the lack of correlation of differential expression on RNA and protein level. This delay is presumably even longer in the secretome as proteins need to be first produced and subsequently secreted^81^.

We focused our study on the secretome to reduce the complexity of the protein mixture analyzed, and to obtain better peptide coverage of the secreted proteins that play an important role in immune signaling. However, this limited our view on the complete proteome affected by immune stimuli. Also, there is the added complication that only a small number of peptides exist that could discriminate between proteoforms. To detect the proteoforms derived from our long-read sequencing data, much deeper shotgun proteomics must be performed^82^. These limitations are reasons why no evidence of novel transcripts could be validated with the proteome.

Multi-omics approaches are a promising method to further our understanding of immune responses. Our study scratches the surface of biological insight to be reaped from a combination of multi-omics and long-read sequencing data and was hindered only by the aforementioned limitations in the samples themselves. Removing these limitations will undoubtedly result in deeper mechanistic understanding and will translate into better outcomes for patients. Insights gained from this methodology can be used immediately in rare disease diagnostics applications, such as the reannotation of variants using more accurate reference transcriptomes for specific tissues^83^, contributing to the development of more personalized medicine.

## Acknowledgements

### Funding

MGN was supported by an ERC Advanced Grant (#833247) and a Spinoza Grant of the Netherlands Organization for Scientific Research. AH was supported by the Solve-RD project, which has received funding from the European Union’s Horizon 2020 research and innovation programme under grant agreement No. 779257.

## Acknowledgement

None

## Author contributions

Conceptualization: RS, PJV, AH, PACH; data curation: RS; formal analysis: RS; investigation: RS, EEV, CIM, TVR, TH; resources: SK, MS; software: RS, TVR; supervision: MM, MGN, PJV, AH, PACH; visualization: RS, EEV; writing—original draft: RS, EEV; writing—review and editing: All authors. All authors read and approved the final manuscript.

## Declaration of interests

### Competing interests

The authors declare no competing interests.

### Consent for publication

Not applicable

### Ethics approval and consent to participate

PBMCs were retrieved form healthy, anonymized donors, as part of the human functional genomics project (HFGP). The HFGP study was approved by the Ethical Committee of Radboud University Nijmegen, the Netherlands (no. 42561.091.12). Experiments were conducted according to the principles expressed in the Declaration of Helsinki. Samples of venous blood were drawn after informed consent was obtained.

### Availability of data and material

Raw PacBio sequencing data and transcriptome is available on EGA under accession number EGAS00001006779. Raw QuantSeq sequencing data is available on EGA under accession number EGAS50000000007. The mass spectrometry proteomics data have been deposited to the ProteomeXchange Consortium via the PRIDE^84^ partner repository with the dataset identifier PXD045237 and 10.6019/PXD045237. Scripts used to generate the results described in this paper can be found at https://github.com/cmbi/hpi_isoseq_paper.

## Notes

https://ega-archive.org/studies/EGAS00001006779

https://github.com/cmbi/hpi_isoseq_paper

### Competing Interest Statement

The authors have declared no competing interest.

### Summary of Updates

Added additional examples of pathogen-specific isoform switching in the manuscript. We describe pathogen specific isoform expression in OAS1 and CLEC7A, along with a corresponding supplementary figure.

## References

1 Medzhitov R, Horng T. Transcriptional control of the inflammatory response. Nature Reviews Immunology 2009 9:10 2009; 9: 692–703.

2 Carpenter S, Ricci EP, Mercier BC, Moore MJ, Fitzgerald KA. Post-transcriptional regulation of gene expression in innate immunity. Nature Reviews Immunology 2014 14:6 2014; 14: 361–376.

3 Wells CA, Chalk AM, Forrest A et al. Alternate transcription of the Toll-like receptor signaling cascade. Genome Biol 2006; 7: 1–17.

4 Oosting M, Kerstholt M, ter Horst R et al. Functional and Genomic Architecture of Borrelia burgdorferi-Induced Cytokine Responses in Humans. Cell Host Microbe 2016; 20: 822–833.

5 Li Y, Oosting M, Smeekens SP et al. A Functional Genomics Approach to Understand Variation in Cytokine Production in Humans. Cell 2016; 167: 1099–1110.e14.

6 Lu YC, Yeh WC, Ohashi PS. LPS/TLR4 signal transduction pathway. Cytokine 2008; 42: 145–151.

7 Alexopoulou L, Holt AC, Medzhitov R, Flavell RA. Recognition of double-stranded RNA and activation of NF-kappaB by Toll-like receptor 3. Nature 2001; 413: 732–738.

8 Van Der Made CI, Simons A, Schuurs-Hoeijmakers J et al. Presence of Genetic Variants Among Young Men With Severe COVID-19. JAMA 2020; 324: 663–673.

9 Smeekens SP, Ng A, Kumar V et al. Functional genomics identifies type I interferon pathway as central for host defense against Candida albicans. Nat Commun 2013; 4. doi:10.1038/NCOMMS2343.

10 Bruno M, Kersten S, Bain JM et al. Transcriptional and functional insights into the host immune response against the emerging fungal pathogen Candida auris. Nat Microbiol 2020; 5: 1516–1531.

11 Askarian F, Wagner T, Johannessen M, Nizet V. Staphylococcus aureus modulation of innate immune responses through Toll-like (TLR), (NOD)-like (NLR) and C-type lectin (CLR) receptors. FEMS Microbiol Rev 2018; 42: 656–671.

12 Netea MG, Gow NAR, Munro CA et al. Immune sensing of Candida albicans requires cooperative recognition of mannans and glucans by lectin and Toll-like receptors. J Clin Invest 2006; 116: 1642–1650.

13 Heinhuis B, Koenders MI, Van De Loo FA, Netea MG, Van Den Berg WB, Joosten LAB. Inflammation-dependent secretion and splicing of IL-32γ in rheumatoid arthritis. Proc Natl Acad Sci U S A 2011; 108: 4962–4967.

14 Oberdoerffer S, Moita LF, Neems D, Freitas RP, Hacohen N, Rao A. Regulation of CD45 alternative splicing by heterogeneous ribonucleoprotein, hnRNPLL. Science 2008; 321: 686–691.

15 Sharon D, Tilgner H, Grubert F, Snyder M. A single-molecule long-read survey of the human transcriptome. Nat Biotechnol 2013; 31: 1009–1014.

16 Glinos DA, Garborcauskas G, Hoffman P et al. Transcriptome variation in human tissues revealed by long-read sequencing. Nature 2022 608:7922 2022; 608: 353–359.

17 Al’Khafaji AM, Smith JT, Garimella K V. et al. High-throughput RNA isoform sequencing using programmed cDNA concatenation. Nature Biotechnology 2023 2023; : 1–5.

18 Vollmers AC, Mekonen HE, Campos S, Carpenter S, Vollmers C. Generation of an isoform-level transcriptome atlas of macrophage activation. J Biol Chem 2021; 296. doi:10.1016/J.JBC.2021.100784.

19 Cole C, Byrne A, Adams M, Volden R, Vollmers C. Complete characterization of the human immune cell transcriptome using accurate full-length cDNA sequencing. Genome Res 2020; 30: 589–601.

20 Inamo J, Suzuki A, Ueda M et al. Immune Isoform Atlas: Landscape of alternative splicing in human immune cells. bioRxiv 2022; : 2022.09.13.507708.

21 Kanno T, Konno R, Miyako K et al. Characterization of proteogenomic signatures of differentiation of CD4+ T cell subsets. DNA Research 2023; 30: 1–11.

22 Shi ZR, Duan YX, Cui F et al. Integrated proteogenomic characterization reveals an imbalanced hepatocellular carcinoma microenvironment after incomplete radiofrequency ablation. Journal of Experimental and Clinical Cancer Research 2023; 42: 1–18.

23 Proteogenomics R, Bautista De Sanctis J, Subbannayya Y et al. Proteogenomics Analysis Reveals Novel Micropeptides in Primary Human Immune Cells. Immuno 2022*, Vol* 2, *Pages 283-292* 2022; **2**: 283–292.

24 Rivero-Hinojosa S, Grant M, Panigrahi A et al. Proteogenomic discovery of neoantigens facilitates personalized multi-antigen targeted T cell immunotherapy for brain tumors. Nature Communications 2021 12:1 2021; 12: 1–15.

25 Oosting M, Kerstholt M, ter Horst R et al. Functional and Genomic Architecture of Borrelia burgdorferi-Induced Cytokine Responses in Humans. Cell Host Microbe 2016; 20: 822–833.

26 Prjibelski A, Mikheenko A, Joglekar A, Smetanin A, Lapidus A, Tilgner H. IsoQuant: a tool for accurate novel isoform discovery with long reads. 2022. doi:10.21203/RS.3.RS-1571850/V1.

27 Martin M. Cutadapt removes adapter sequences from high-throughput sequencing reads. EMBnet J 2011; 17: 10–12.

28 Patro R, Duggal G, Love MI, Irizarry RA, Kingsford C. Salmon provides fast and bias-aware quantification of transcript expression. Nat Methods 2017; 14: 417–419.

29 Tarazona S, Furió-Tarí P, Turrà D et al. Data quality aware analysis of differential expression in RNA-seq with NOISeq R/Bioc package. Nucleic Acids Res 2015; 43: e140.

30 Ashburner M, Ball CA, Blake JA et al. Gene Ontology: tool for the unification of biology. Nature Genetics 2000 25:1 2000; 25: 25–29.

31 Wingender E, Dietze P, Karas H, Knüppel R. TRANSFAC: a database on transcription factors and their DNA binding sites. Nucleic Acids Res 1996; 24: 238–241.

32 Reese F, Mortazavi A. Swan: a library for the analysis and visualization of long-read transcriptomes. Bioinformatics 2021; 37: 1322–1323.

33 Vitting-Seerup K, Sandelin A, Berger B. IsoformSwitchAnalyzeR: analysis of changes in genome-wide patterns of alternative splicing and its functional consequences. Bioinformatics 2019; 35: 4469–4471.

34 Wang L, Park HJ, Dasari S, Wang S, Kocher JP, Li W. CPAT: coding-Potential Assessment Tool using an alignment-free logistic regression model. Nucl Acids Res 2013; 41: e74.

35 Eddy SR. HMMER User’s Guide Biological sequence analysis using profile hidden Markov models. 2020.http://hmmer.org (accessed 2 Feb2022).

36 Almagro Armenteros JJ, Tsirigos KD, Sønderby CK et al. SignalP 5.0 improves signal peptide predictions using deep neural networks. Nature Biotechnology 2019 37:4 2019; 37: 420–423.

37 Shannon P, Markiel A, Ozier O et al. Cytoscape: a software environment for integrated models of biomolecular interaction networks. Genome Res 2003; 13: 2498–2504.

38 Merico D, Isserlin R, Stueker O, Emili A, Bader GD. Enrichment Map: A Network-Based Method for Gene-Set Enrichment Visualization and Interpretation. PLoS One 2010; 5: e13984.

39 Fang H. dcGOR: an R package for analysing ontologies and protein domain annotations. PLoS Comput Biol 2014; 10. doi:10.1371/JOURNAL.PCBI.1003929.

40 Miller RM, Jordan BT, Mehlferber MM et al. Enhanced protein isoform characterization through long-read proteogenomics. Genome Biol 2022; 23. doi:10.1186/s13059-022-02624-y.

41 Miller RM, Millikin RJ, Rolfs Z, Shortreed MR, Smith LM. Enhanced Proteomic Data Analysis with MetaMorpheus. Methods Mol Biol 2023; 2426: 35–66.

42 Deutsch EW, Lane L, Overall CM et al. Human Proteome Project Mass Spectrometry Data Interpretation Guidelines 3.0. J Proteome Res 2019; 18: 4108–4116.

43 Millikin RJ, Shortreed MR, Scalf M, Smith LM. Fast, Free, and Flexible Peptide and Protein Quantification with FlashLFQ. Methods Mol Biol 2023; 2426: 303–313.

44 Digre A, Lindskog C. The Human Protein Atlas-Spatial localization of the human proteome in health and disease. Protein Sci 2021; 30: 218–233.

45 Gu Z, Eils R, Schlesner M. Complex heatmaps reveal patterns and correlations in multidimensional genomic data. Bioinformatics 2016; 32: 2847–2849.

46 Noguchi S, Arakawa T, Fukuda S et al. FANTOM5 CAGE profiles of human and mouse samples. Sci Data 2017; 4. doi:10.1038/SDATA.2017.112.

47 Moll P, Ante M, Seitz A, Reda T. QuantSeq 3ʹ mRNA sequencing for RNA quantification. Nature Methods 2014 11:12 2014; 11: i–iii.

48 Raudvere U, Kolberg L, Kuzmin I et al. g:Profiler: a web server for functional enrichment analysis and conversions of gene lists (2019 update). Nucleic Acids Res 2019; 47: W191–W198.

49 Paun A, Pitha PM. The IRF family, revisited. Biochimie 2007; 89: 744–753.

50 Herbein G, Doyle AG, Montaner LJ, Gordon S. Lipopolysaccharide (LPS) down-regulates CD4 expression in primary human macrophages through induction of endogenous tumour necrosis factor (TNF) and IL-1 beta. Clin Exp Immunol 1995; 102: 430–437.

51 Lachmandas E, Boutens L, Ratter JM et al. Microbial stimulation of different Toll-like receptor signalling pathways induces diverse metabolic programmes in human monocytes. Nat Microbiol 2016; 2. doi:10.1038/NMICROBIOL.2016.246.

52 Green ID, Pinello N, Song R et al. Macrophage development and activation involve coordinated intron retention in key inflammatory regulators. Nucleic Acids Res 2020; 48: 6513–6529.

53 Song R, Tikoo S, Jain R et al. Dynamic intron retention modulates gene expression in the monocytic differentiation pathway. Immunology 2022; 165: 274–286.

54 Wong JJL, Ritchie W, Ebner OA et al. Orchestrated intron retention regulates normal granulocyte differentiation. Cell 2013; 154. doi:10.1016/J.CELL.2013.06.052.

55 Ullrich S, Guigó R. Dynamic changes in intron retention are tightly associated with regulation of splicing factors and proliferative activity during B-cell development. Nucleic Acids Res 2020; 48: 1327–1340.

56 O’Grady TM, Baddoo M, Flemington SA, Ishaq EY, Ungerleider NA, Flemington EK. Reversal of splicing infidelity is a pre-activation step in B cell differentiation. Front Immunol 2022; 13. doi:10.3389/FIMMU.2022.1060114.

57 Karginov TA, Ménoret A, Vella AT. Optimal CD8+ T cell effector function requires costimulation-induced RNA-binding proteins that reprogram the transcript isoform landscape. Nat Commun 2022; 13. doi:10.1038/S41467-022-31228-0.

58 Ni T, Yang W, Han M et al. Global intron retention mediated gene regulation during CD4+ T cell activation. Nucleic Acids Res 2016; 44: 6817–6829.

59 Brunet MA, Levesque SA, Hunting DJ, Cohen AA, Roucou X. Recognition of the polycistronic nature of human genes is critical to understanding the genotype-phenotype relationship. Genome Res 2018; 28: 609–624.

60 Heinz S, Texari L, Hayes MGB et al. Transcription Elongation Can Affect Genome 3D Structure. Cell 2018; 174: 1522–1536.e22.

61 Karasawa T, Kawashima A, Usui F et al. Oligomerized CARD16 promotes caspase-1 assembly and IL-1β processing. FEBS Open Bio 2015; 5: 348–356.

62 Devi S, Indramohan M, Jäger E et al. CARD-only proteins regulate in vivo inflammasome responses and ameliorate gout. Cell Rep 2023; 42. doi:10.1016/J.CELREP.2023.112265.

63 Oeckinghaus A, Ghosh S. The NF-kappaB family of transcription factors and its regulation. Cold Spring Harb Perspect Biol 2009; 1. doi:10.1101/CSHPERSPECT.A000034.

64 Fliegauf M, Kinnunen M, Posadas-Cantera S et al. Detrimental NFKB1 missense variants affecting the Rel-homology domain of p105/p50. Front Immunol 2022; 13. doi:10.3389/FIMMU.2022.965326.

65 Mata-Martínez P, Bergón-Gutiérrez M, del Fresno C. Dectin-1 Signaling Update: New Perspectives for Trained Immunity. Front Immunol 2022; 13: 812148.

66 Rotival M, Quach H, Quintana-Murci L. Defining the genetic and evolutionary architecture of alternative splicing in response to infection. Nat Commun 2019; 10: 1– 15.

67 Banday AR, Stanifer ML, Florez-Vargas O et al. Genetic regulation of OAS1 nonsense-mediated decay underlies association with COVID-19 hospitalization in patients of European and African ancestries. Nature Genetics 2022 54:8 2022; 54: 1103–1116.

68 Wickenhagen A, Sugrue E, Lytras S et al. A prenylated dsRNA sensor protects against severe COVID-19. Science (1979) 2021; 374. doi:10.1126/SCIENCE.ABJ3624/SUPPL_FILE/SCIENCE.ABJ3624_MDAR_REPRODUCIBILITY_CHECKLIST.PDF.

69 Perišić Nanut M, Pečar Fonović U, Jakoš T, Kos J. The Role of Cysteine Peptidases in Hematopoietic Stem Cell Differentiation and Modulation of Immune System Function. Front Immunol 2021; 12. doi:10.3389/FIMMU.2021.680279.

70 Thiery J, Keefe D, Saffarian S et al. Perforin activates clathrin- and dynamin-dependent endocytosis, which is required for plasma membrane repair and delivery of granzyme B for granzyme-mediated apoptosis. Blood 2010; 115: 1582–1593.

71 Momoi T, Suzuki M, Titani K, Hisanaga S, Ogawa H, Saito A. Amino acid sequence of a modified β2-microglobulin in renal failure patient urine and long-term dialysis patient blood. Clinica Chimica Acta 1995; 236: 135–144.

72 Fukuhara K, Okumura M, Shiono H et al. A study on CD45 isoform expression during T-cell development and selection events in the human thymus. Hum Immunol 2002; 63: 394–404.

73 Orta-Mascaró M, Consuegra-Fernández M, Carreras E et al. CD6 modulates thymocyte selection and peripheral T cell homeostasis. J Exp Med 2016; 213: 1387– 1397.

74 De Arras L, Alper S. Limiting of the innate immune response by SF3A-dependent control of MyD88 alternative mRNA splicing. PLoS Genet 2013; 9. doi:10.1371/JOURNAL.PGEN.1003855.

75 Pozzi B, Bragado L, Mammi P et al. Dengue virus targets RBM10 deregulating host cell splicing and innate immune response. Nucleic Acids Res 2020; 48: 6824–6838.

76 Stein MM, Conery M, Magnaye KM et al. Sex-specific differences in peripheral blood leukocyte transcriptional response to LPS are enriched for HLA region and X chromosome genes. Sci Rep 2021; 11. doi:10.1038/S41598-020-80145-Z.

77 Piasecka B, Duffy D, Urrutia A et al. Distinctive roles of age, sex, and genetics in shaping transcriptional variation of human immune responses to microbial challenges. Proc Natl Acad Sci U S A 2018; 115: E488–E497.

78 Li Y, Oosting M, Deelen P et al. Inter-individual variability and genetic influences on cytokine responses to bacteria and fungi. Nature Medicine 2016 22:8 2016; 22: 952–960.

79 Mironov A, Petrova M, Margasyuk S et al. Tissue-specific regulation of gene expression via unproductive splicing. Nucleic Acids Res 2023; 51: 3055–3066.

80 Hardwick SA, Hu W, Joglekar A et al. Single-nuclei isoform RNA sequencing unlocks barcoded exon connectivity in frozen brain tissue. Nat Biotechnol 2022; 40: 1082– 1092.

81 Meissner F, Scheltema RA, Mollenkopf HJ, Mann M. Direct proteomic quantification of the secretome of activated immune cells. Science (1979) 2013; 340: 475–478.

82 Sinitcyn P, Richards AL, Weatheritt RJ et al. Global detection of human variants and isoforms by deep proteome sequencing. Nature Biotechnology 2023 2023; : 1–11.

83 Salz R, Saraiva-Agostinho N, Vorsteveld E et al. SUsPECT: a pipeline for variant effect prediction based on custom long-read transcriptomes for improved clinical variant annotation. BMC Genomics 2023; 24: 1–10.

84 Perez-Riverol Y, Bai J, Bandla C et al. The PRIDE database resources in 2022: a hub for mass spectrometry-based proteomics evidences. Nucleic Acids Res 2022; 50: D543– D552.

