## Supplementary figures for "Multi-omic profiling of pathogen-stimulated primary immune cells"

### Slide 1
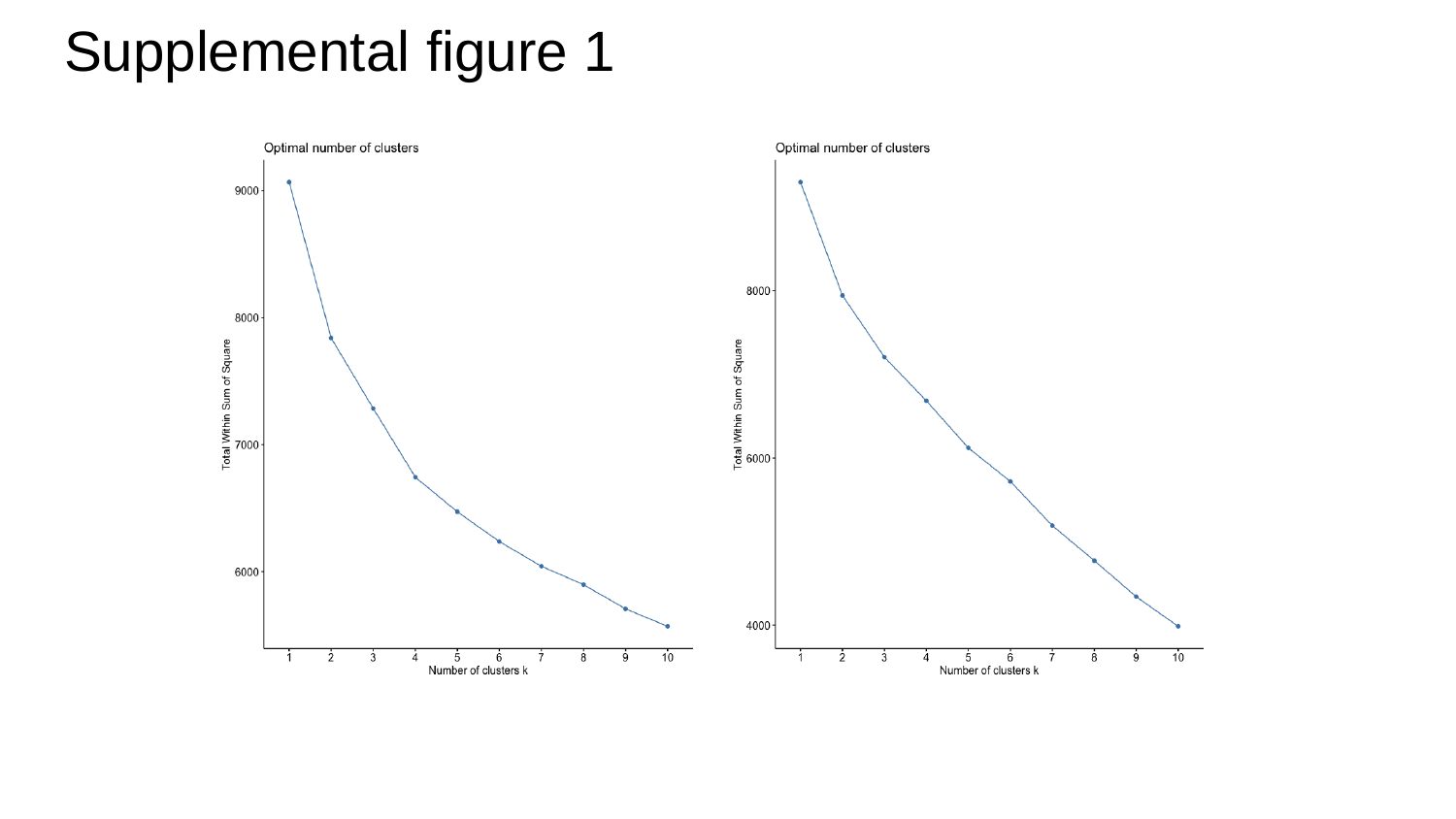

#### Supplemental figure 1

### Slide 2
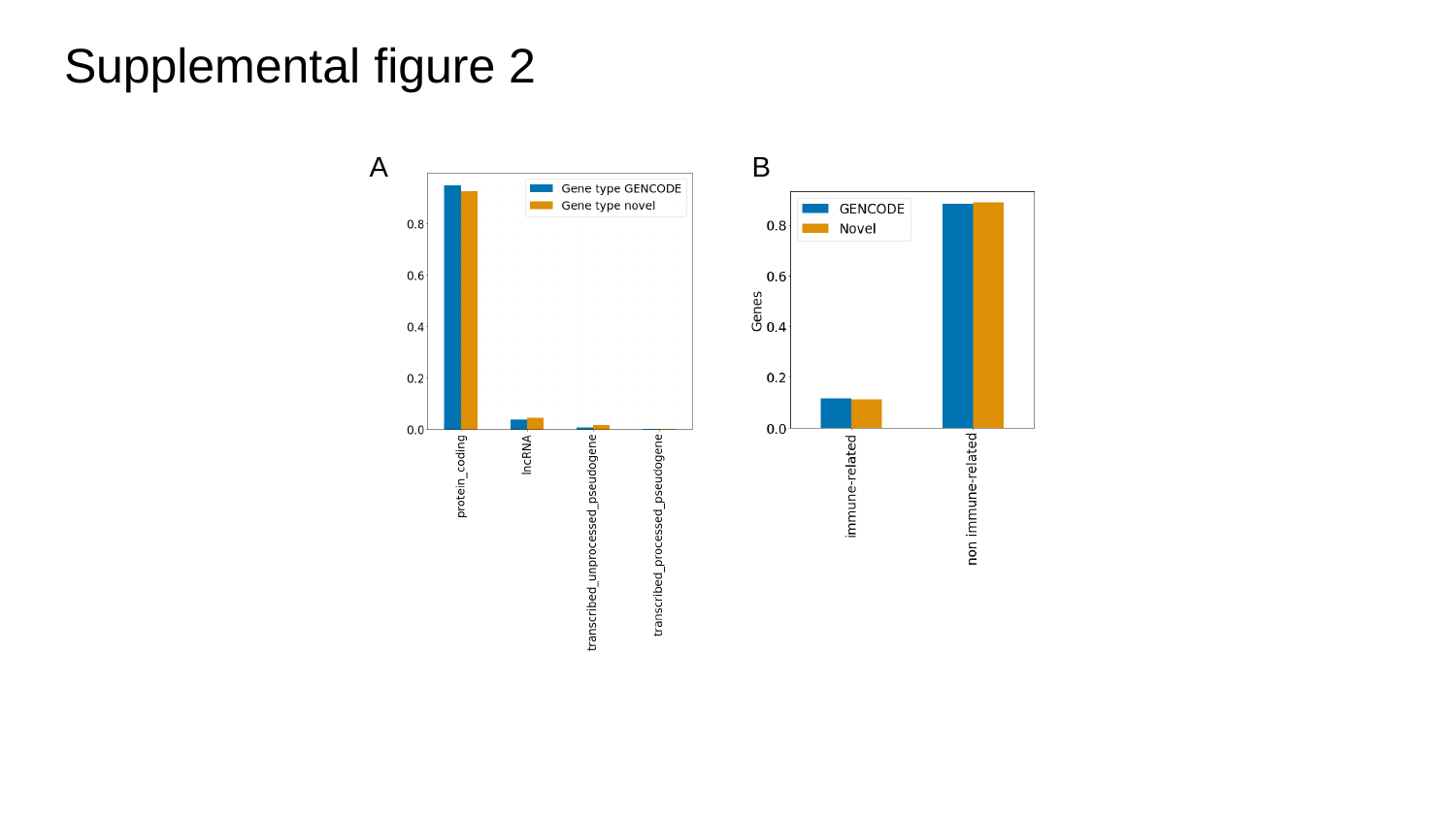

#### Supplemental figure 2
A
B

### Slide 3
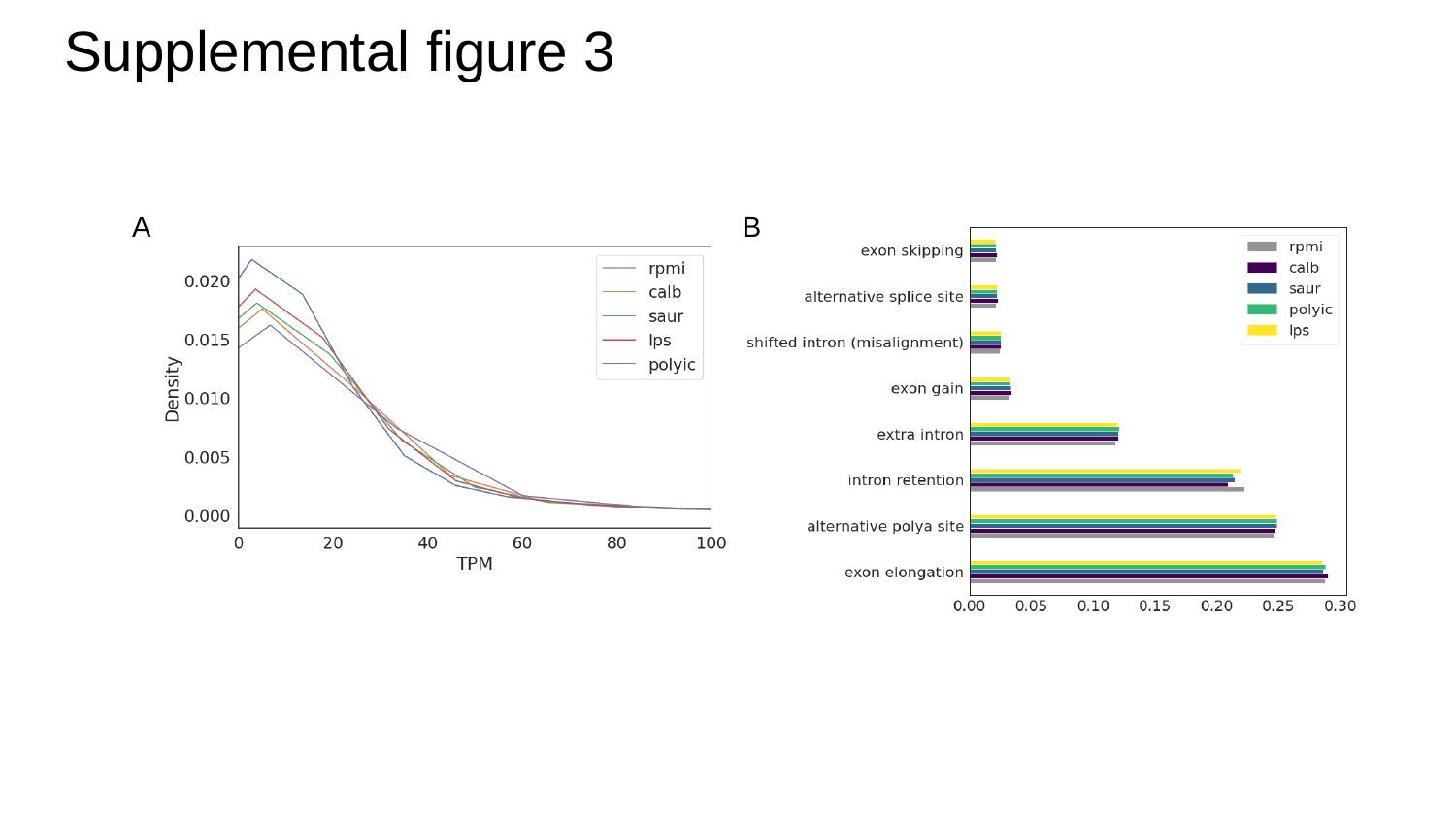

#### Supplemental figure 3
A
B

### Slide 4
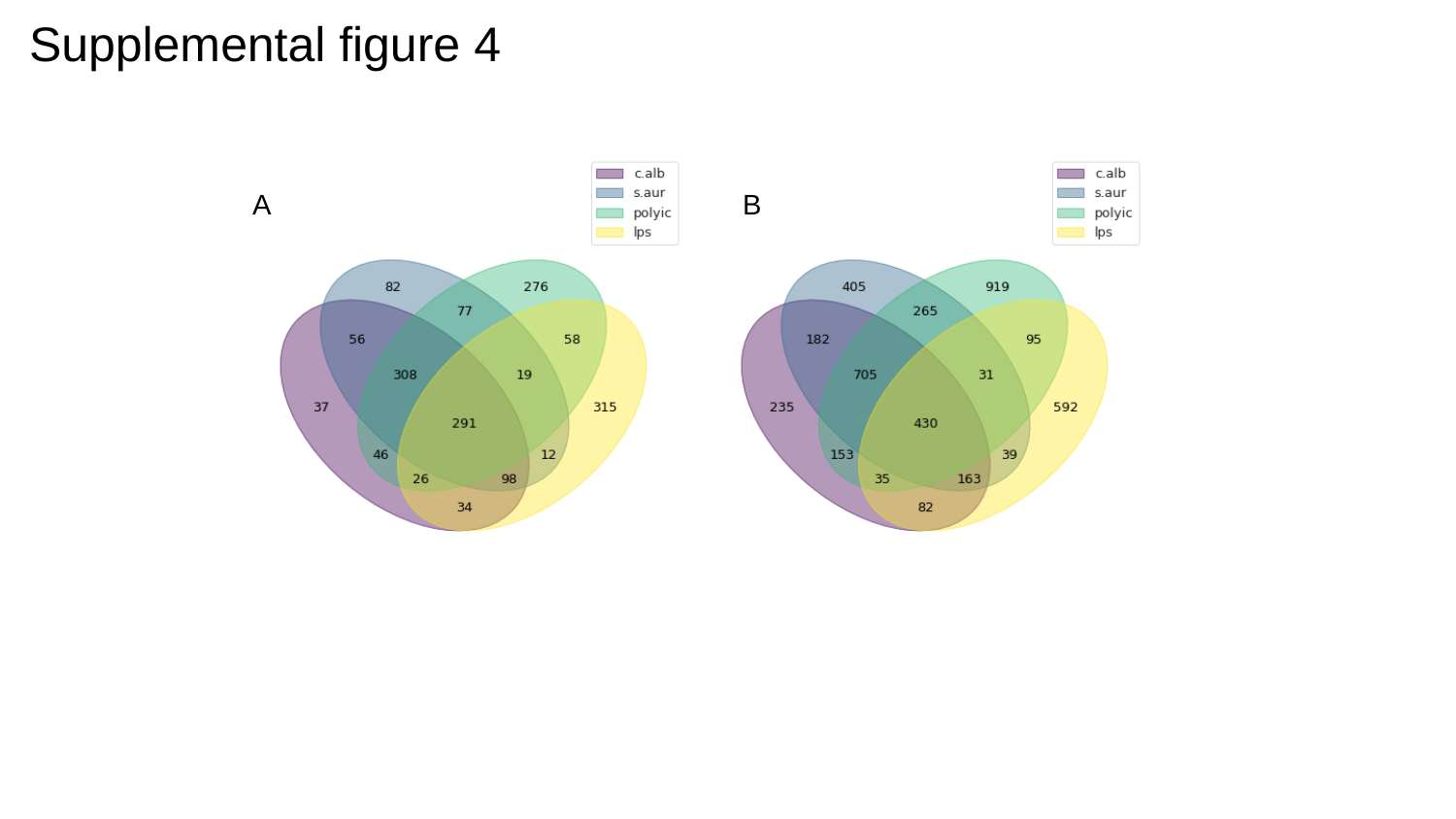

#### Supplemental figure 4
A
B

### Slide 5
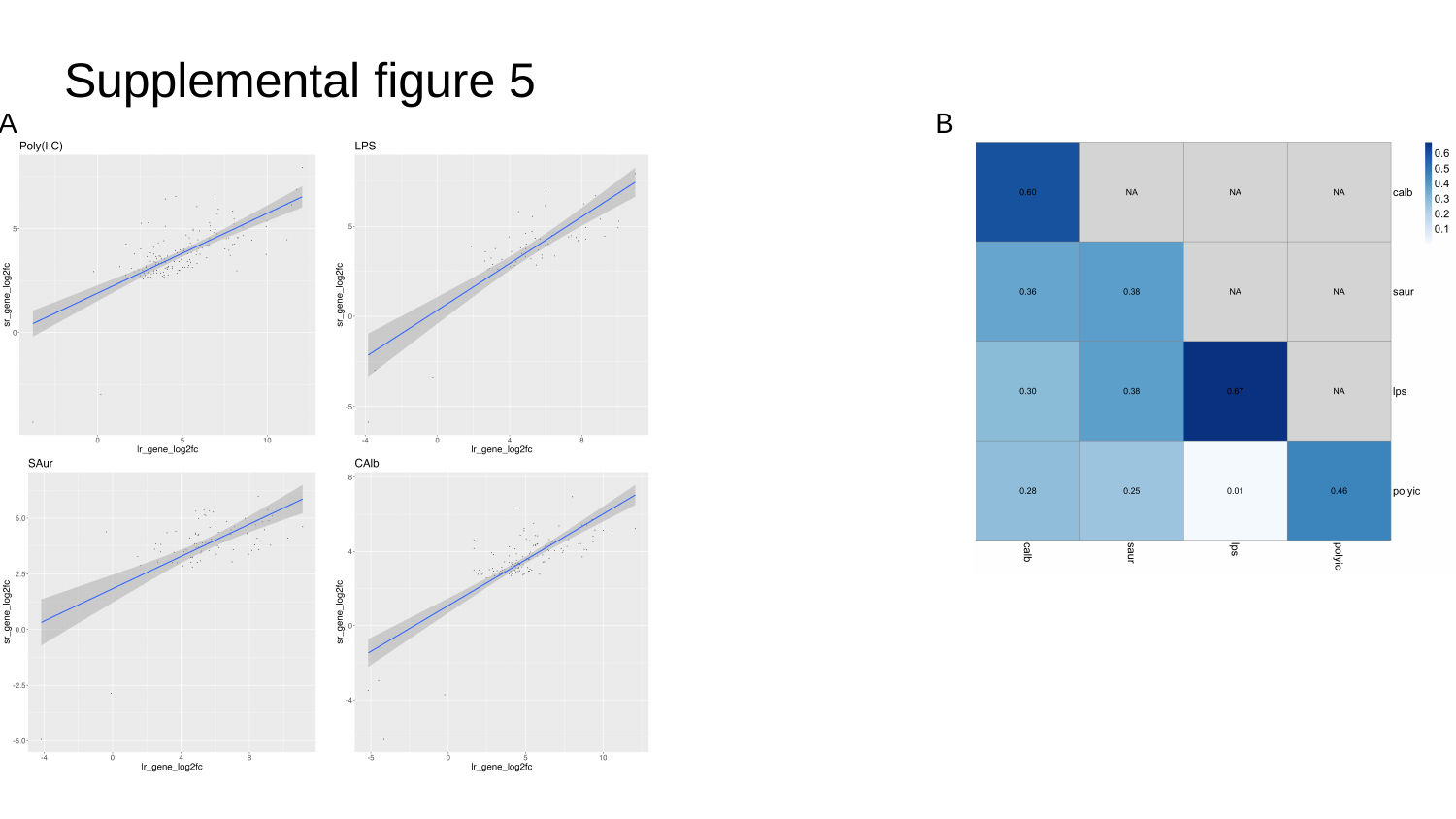

#### Supplemental figure 5
A
B

### Slide 6
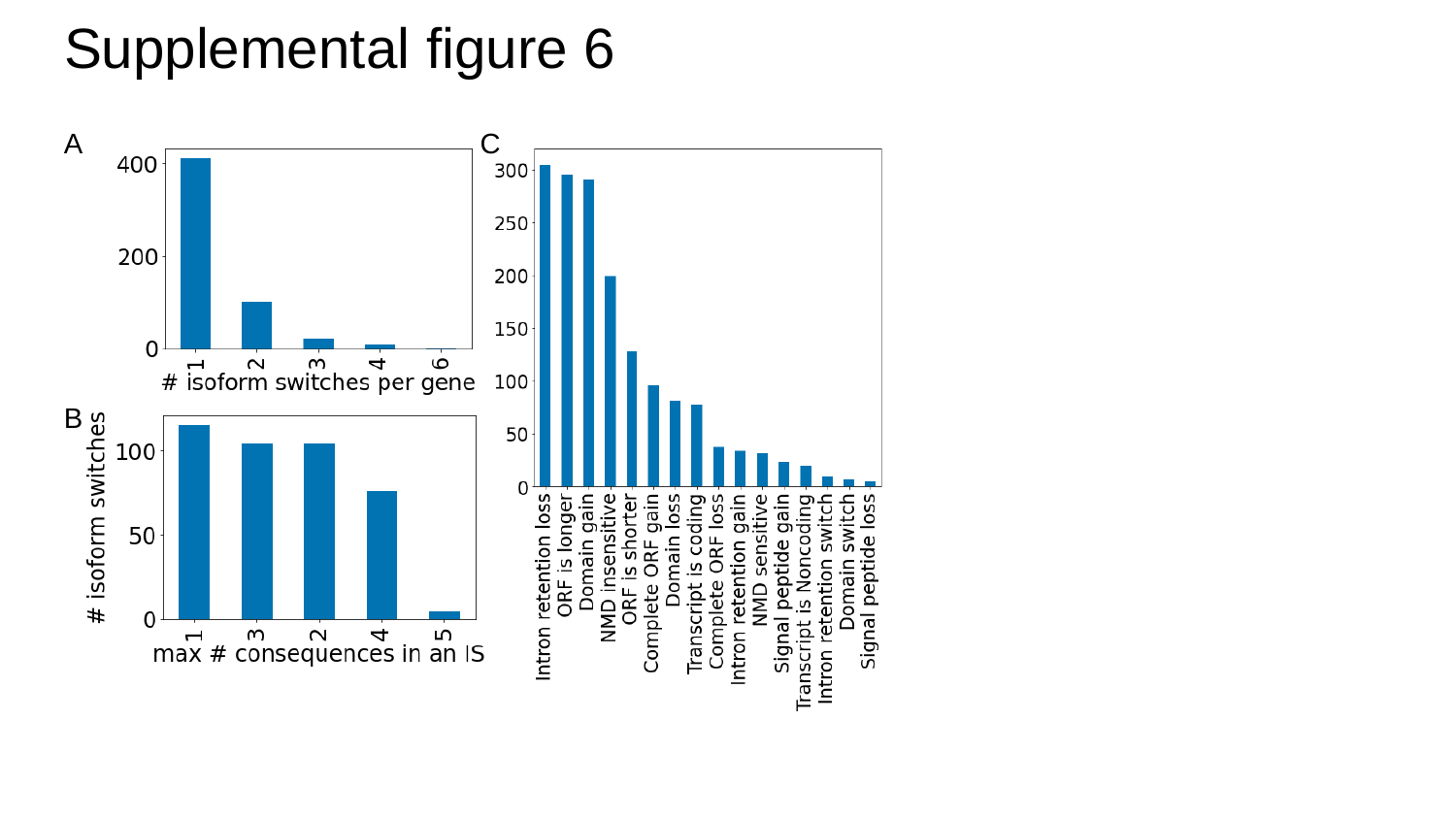

#### Supplemental figure 6
C
A
B

### Slide 7
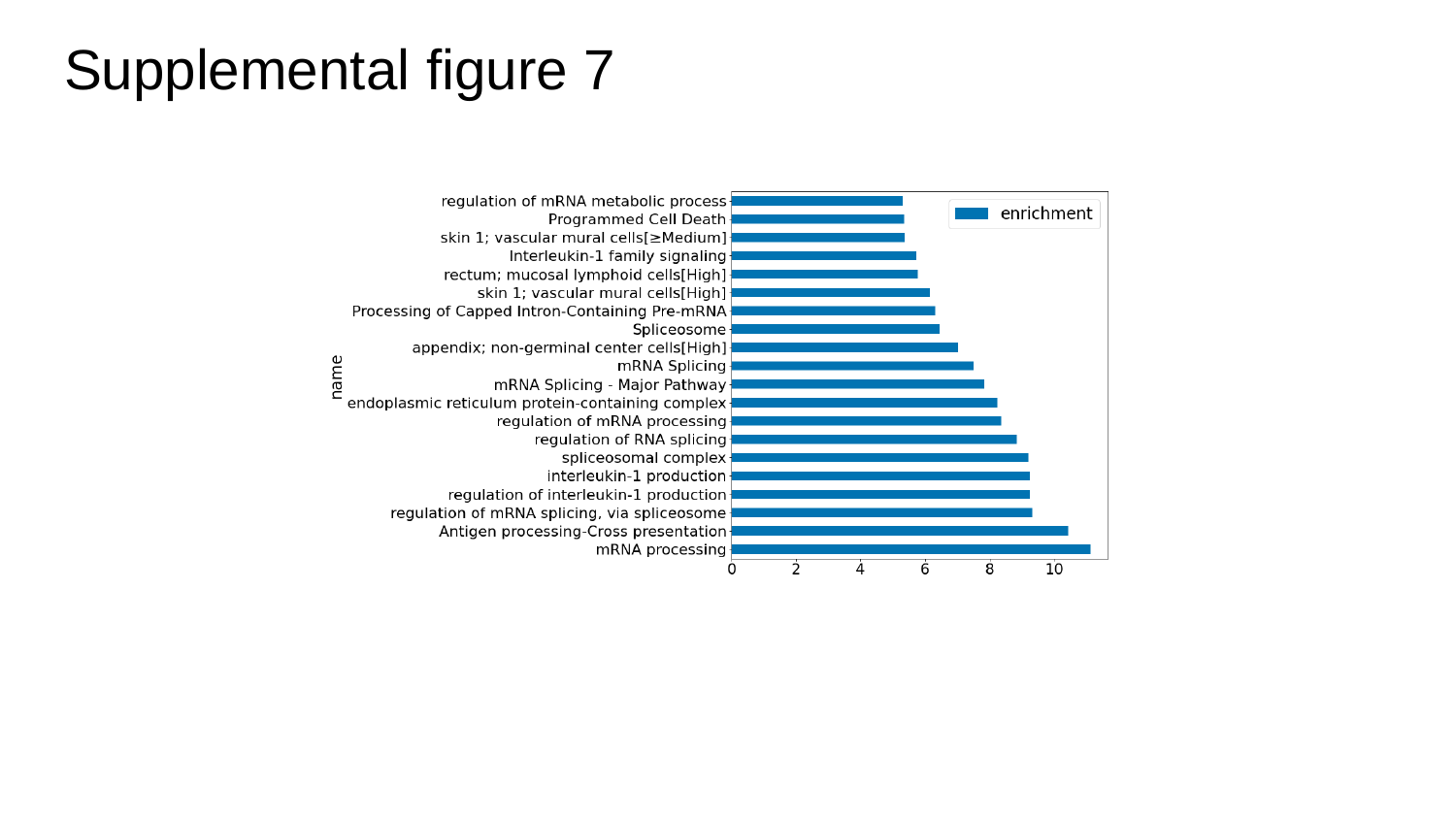

Supplemental figure 7

### Slide 8
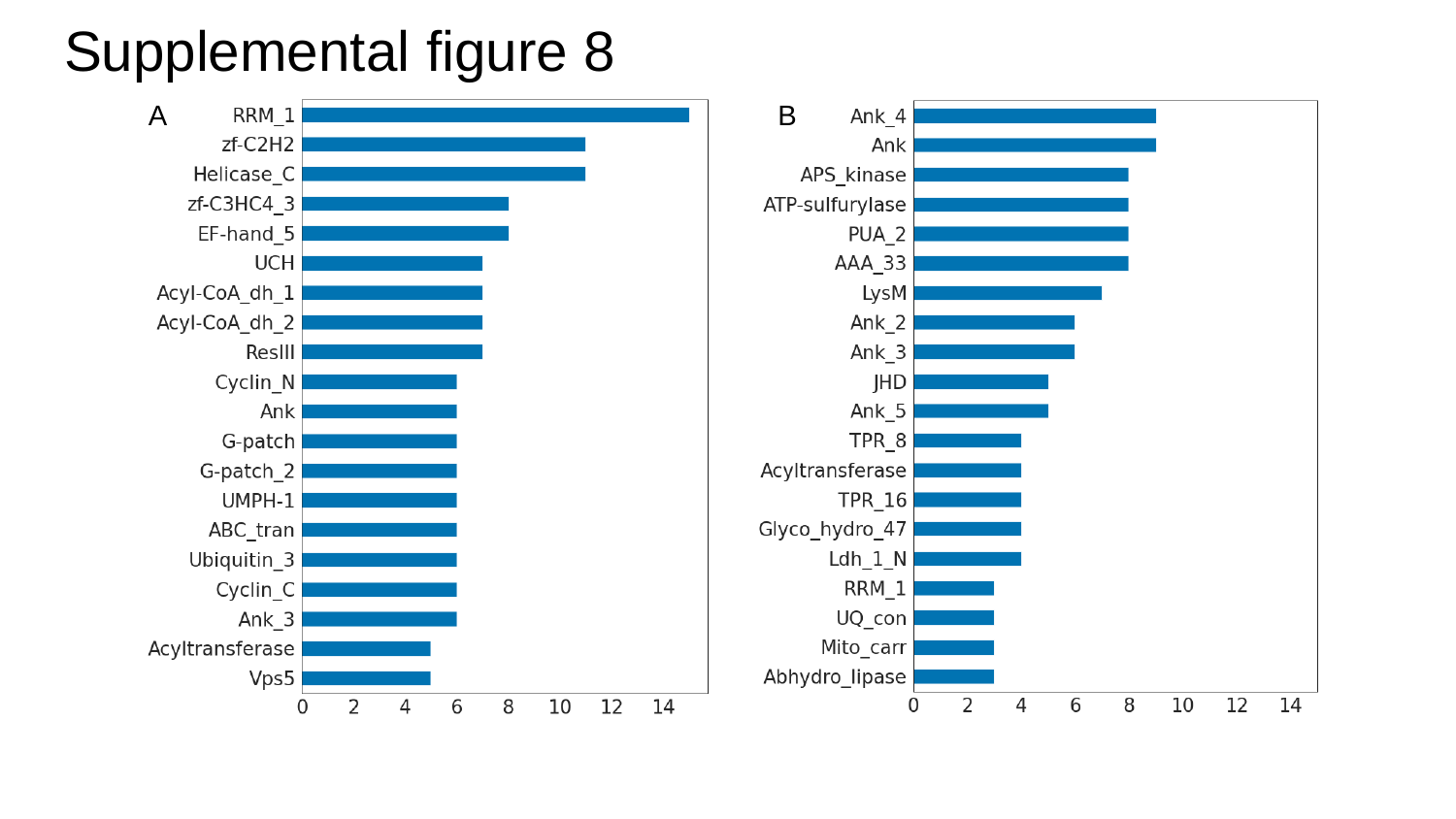

#### Supplemental figure 8
B
A

### Slide 9
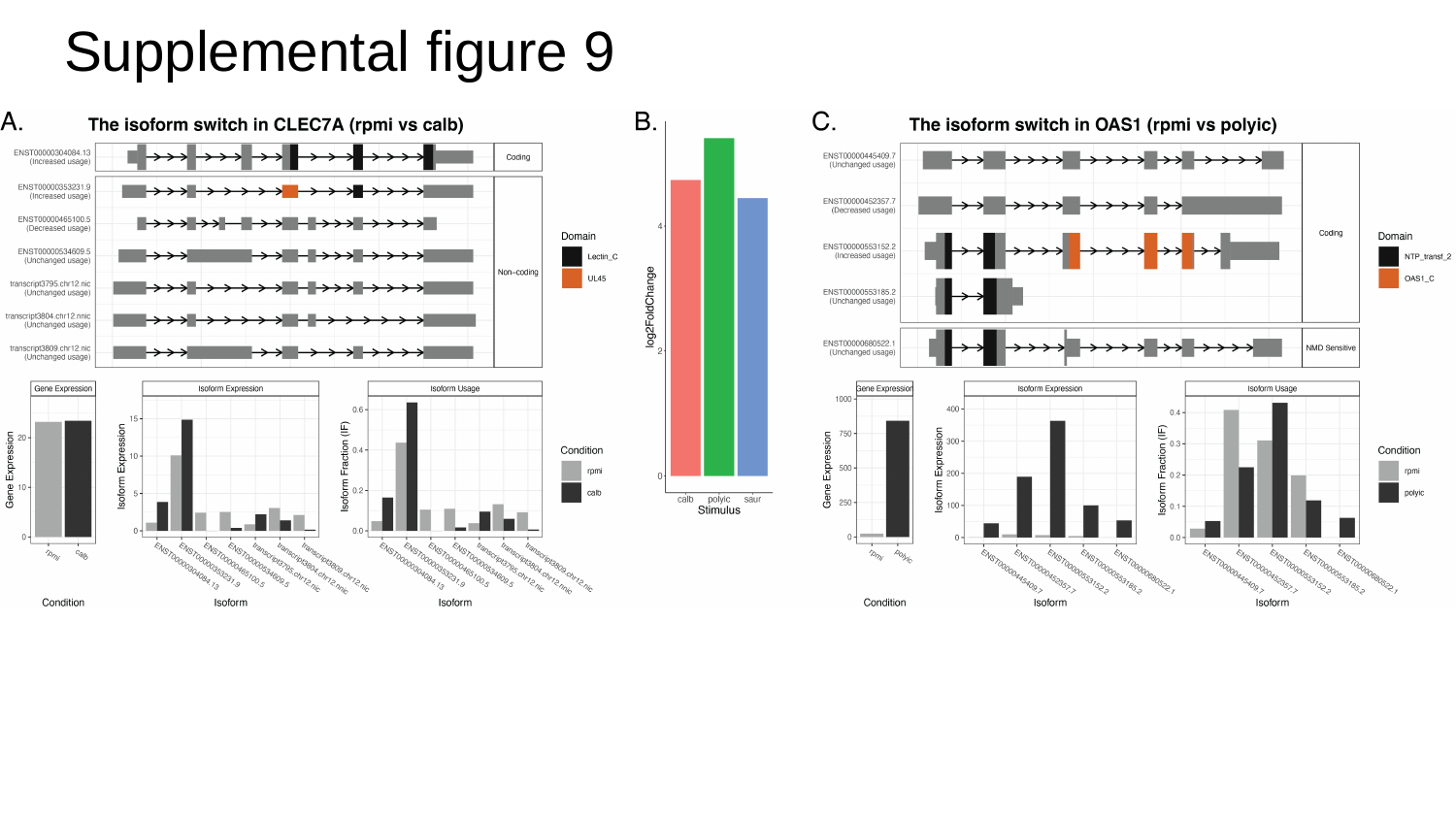

#### Supplemental figure 9

### Slide 10
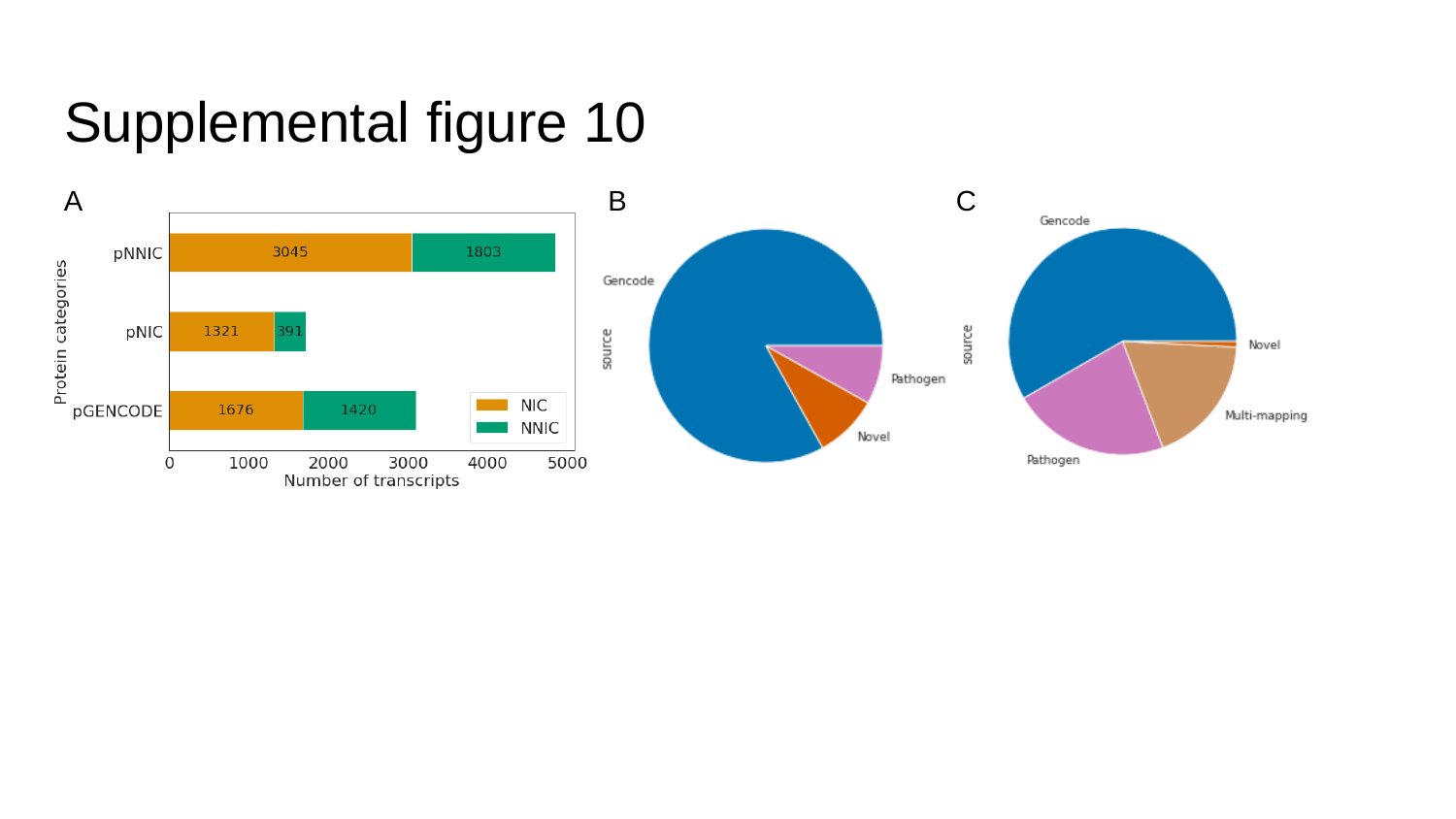

#### Supplemental figure 10
A
B
C

### Slide 11
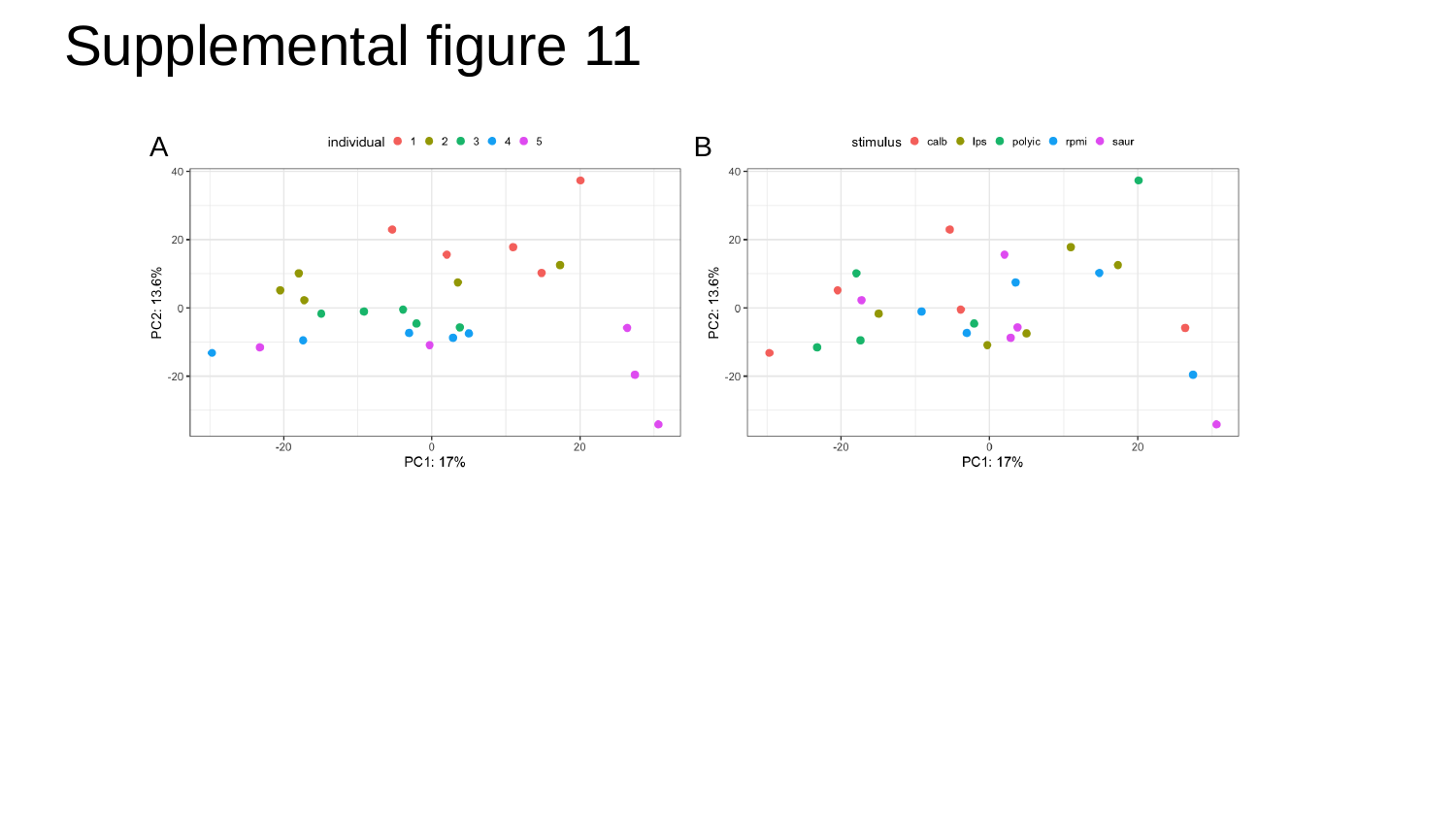

#### Supplemental figure 11
A
B

### Slide 12
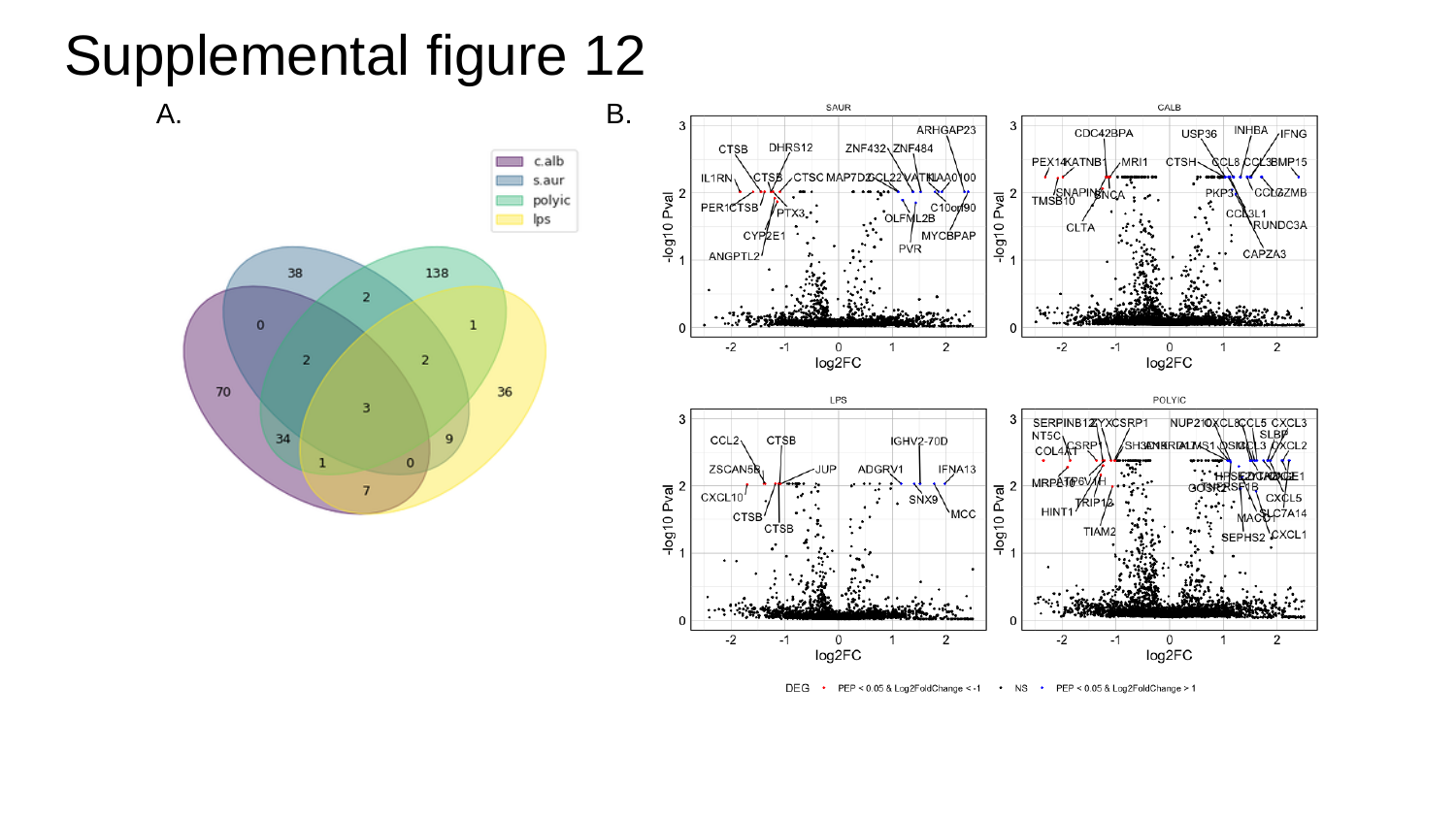

#### Supplemental figure 12
A.
B.
